# tRNA anticodon cleavage by target-activated CRISPR-Cas13a effector

**DOI:** 10.1101/2021.11.10.468108

**Authors:** Ishita Jain, Matvey Kolesnik, Leonid Minakhin, Natalia Morozova, Anna Shiriaeva, Alexandr Kirillov, Sofia Medvedeva, Konstantin Kuznedelov, Sergei Borukhov, Kira S. Makarova, Eugene V. Koonin, Konstantin Severinov, Ekaterina Semenova

**Author notes:** These authors contributed equally: Ishita Jain, Matvey Kolesnik. Present address: Department of Biochemistry and Molecular Biology, Thomas Jefferson University, Philadelphia, USA.

## Abstract

Type VI CRISPR-Cas systems are the only CRISPR variety that cleaves exclusively RNA^1,2^. In addition to the CRISPR RNA (crRNA)-guided, sequence-specific binding and cleavage of target RNAs, such as phage transcripts, the type VI effector, Cas13, causes collateral RNA cleavage, which induces bacterial cell dormancy, thus protecting the host population from phage spread^3,4^. We show here that the principal form of collateral RNA degradation elicited by Cas13a protein from *Leptotrichia shahii* upon target RNA recognition is the cleavage of anticodons of multiple tRNA species, primarily those with anticodons containing uridines. This tRNA cleavage is necessary and sufficient for bacterial dormancy induction by Cas13a. In addition, Cas13a activates the RNases of bacterial toxin-antitoxin modules, thus indirectly causing mRNA and rRNA cleavage, which could provide a back-up defense mechanism. The identified mode of action of Cas13a resembles that of bacterial anticodon nucleases involved in antiphage defense^5^, which is compatible with the hypothesis that type VI effectors evolved from an abortive infection module^6,7^ encompassing an anticodon nuclease.

---

Among the numerous, evolutionarily and mechanistically diverse CRISPR-Cas systems, type VI stands out for the exclusive targeting of RNA by the single-subunit effector RNase Cas13^3,8^. Type VI CRISPR-Cas provide efficient protection against RNA phages in laboratory experiments^3^ as well as against DNA phages, the dominant component of bacterial viromes, via phage transcript cleavage^4,9^. Target RNA binding by Cas13 programmed with a cognate crRNA triggers a conformational rearrangement of the two HEPN (*H*igher *E*ukaryotes and *P*rokaryotes *N*ucleotide-binding) domains of Cas13, which form an active nuclease that catalyzes both target RNA cleavage *in cis* and collateral cleavage of non-target RNAs *in trans*^3,10–12^. Collateral RNA cleavage was observed after target recognition *in vitro* and is considered to be non-specific^3,11,12^. In *Escherichia coli*, targeting of non-essential transcripts by Cas13 resulted in cell growth suppression, which was proposed to be a consequence of collateral degradation of essential transcripts^3^. Targeting phage transcripts by *Listeria* Cas13a caused massive degradation of both host and phage RNAs and induced cell dormancy, preventing the spread of phage progeny through the bacterial population^4^.

Presently, it is unclear whether the target-activated Cas13a RNase activity is solely and directly responsible for dormancy induction, and if so, what are the main essential substrates of Cas13 RNase, or it functions by activating other cellular stress response systems. In this work, we investigated the collateral activity of Cas13a from *Leptotrichia shahii* and determined that this protein (LshCas13a), when activated by the recognition of target RNAs, specifically cleaves tRNAs at anticodon loops *in vivo* and *in vitro*. Massive tRNA cleavage by target-activated Cas13a leads to protein synthesis inhibition and cell growth arrest. In addition, Cas13a collateral tRNA cleavage activates cellular RNases encoded by toxin-antitoxin modules. Together, these results reveal the mechanism of type VI CRISPR-Cas immunity, which is found to primarily direct dormancy induction by Cas13a, but also involves the host stress response machinery.

## Cas13a RNA-targeting hinders cell growth by inhibiting protein synthesis

Throughout this work, we use a previously described experimental model of *E. coli* cells transformed with a plasmid containing the *L. shahii* type VI-A CRISPR-Cas system targeting a red fluorescent protein (RFP) mRNA which is inducibly transcribed from a compatible plasmid^3^. We will refer to these cells as “targeting” cells. As a control, “nontargeting” cells containing a type VI-A CRISPR array encoding crRNAs carrying spacers that did not match sequences in the *E. coli* genome or in the plasmids are used in the experiments (Fig. 1a, b). As can be seen from Fig. 1c and in agreement with a previous report^3^, targeting the non-essential RFP transcript substantially decreased the rate of cell growth.

**Fig. 1:**
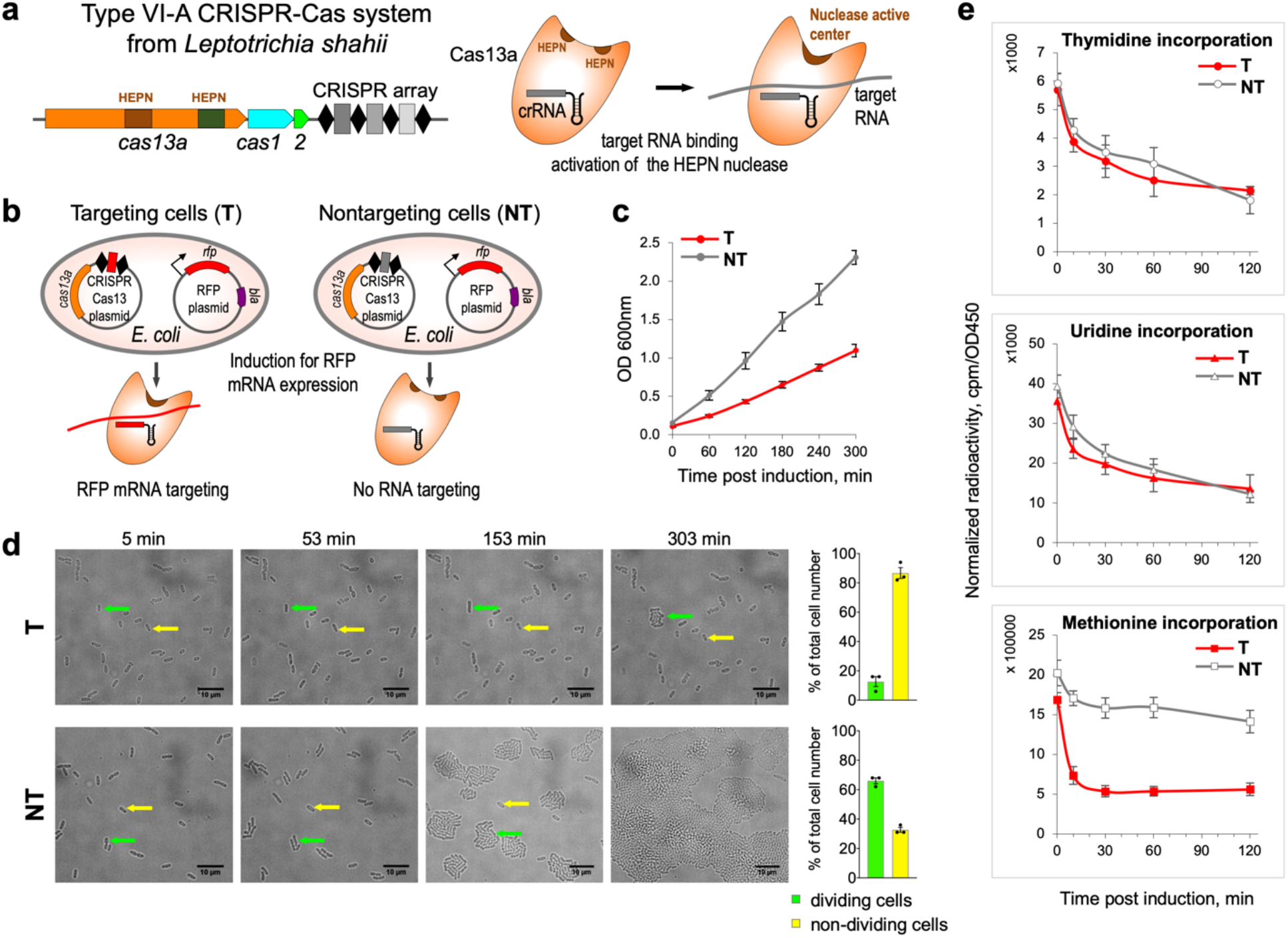
Cell growth arrest coupled with translation inhibition in *E. coli* expressing target-activated LshCas13a. **a**, Schematic of the *Leptotrichia shahii* CRISPR-Cas13a locus encoding Cas13a protein containing the two HEPN domains (brown), Cas1 and Cas2 proteins of adaptation module, and CRISPR array. Target RNA binding to cognate crRNA activates the HEPN nuclease. **b**, RFP mRNA targeting by LshCas13a in *E. coli*. The cells contain CRISPR-Cas13a plasmid that encodes LshCas13a and crRNA, and RFP plasmid that directs inducible expression of RFP and carries the *bla* gene. In targeting cells (T) CRISPR array contains a spacer (shown in red) matching RFP mRNA, in nontargeting control cells (NT) CRISPR array spacer (shown in grey) does not match any RNA in cells. **c**, Targeting of non-essential RFP mRNA by LshCas13a leads to growth suppression in targeting (T, red) cells; no such effect is observed in nontargeting (NT, grey) control. OD_600_ measurements are taken after RFP induction by anhydrotetracycline. OD_600_ values represent a mean ± s. e. m. of three biological replicates at each time point. **d**, Time-lapse microscopy showing dividing (green arrows) and non-dividing (yellow arrows) cells in targeting (T) and non-targeting (NT) samples. *E. coli* cells were grown on LB-agarose blocks containing inducer for RFP expression and antibiotics. Three fields of view for tracking the fate of at least 100 cells were analyzed in each experiment. Bar graph shows mean percentages (± s. e. m.) of dividing and non-dividing cells from the total number of cells counted for targeting (top graph) and nontargeting (bottom graph) samples, obtained from three biological replicates. **e**, Time course showing incorporation of radiolabeled precursors: ^3^H-thymidine, ^3^H-uridine or ^35^S-methionine by pulse-labeling in targeting (T, red) and nontargeting (NT, grey) *E. coli* cells grown in minimal media supplemented with RFP inducer and antibiotics. Radioactivity at each time point is normalized to OD450 of the samples taken after RFP induction. Mean ± s. e. m. for three independent experiments are shown.

Time-lapse microscopy showed that RFP mRNA targeting by the *L. shahii* type VI-A CRISPR-Cas system did not lead to cell lysis or changes in cell morphology, but markedly decreased the proportion of dividing cells in the culture. In contrast, most nontargeting cells divided multiple times during the 300-minute post-induction observation and formed microcolonies (Fig. 1d). Cell viability was tested using membrane-impermeable DNA binding dye YOYO-1 (Extended Data Fig. 1). The membranes of non-dividing targeting cells remained intact and only a small (<2%) fraction of cells were dead 180 minutes post-induction. A similar fraction of dead cells was observed in the nontargeting culture (Extended Data Fig. 1). We thus conclude that targeting a non-essential RNA by Cas13a induces cell dormancy rather than cell death.

To determine which major cellular process(es) was affected by Cas13a RNA targeting, we performed pulse-labeling analysis using radiolabeled precursors of replication, transcription, and translation. ^3^H-thymidine and ^3^H-uridine incorporation were comparable in targeting and nontargeting cells cultures, indicating that target recognition by Cas13a had no major effects on DNA replication or RNA synthesis during transcription. In contrast, ^35^S-methionine incorporation was strongly inhibited in the targeting but not nontargeting cells cultures, indicating that protein synthesis was affected (Fig. 1e). Analysis of radioactively-labeled proteins synthesized post-induction by SDS-PAGE indicated that translation inhibition was not specific to RFP, whose mRNA was targeted by Cas13a. Instead, decreased synthesis of all detectable cellular proteins was apparent as early as 5 minutes post-induction (Extended Data Fig. 2).

## mRNAs, rRNAs, and tRNAs are cleaved at specific sites in targeting cells

To investigate the cause(s) of global translation inhibition observed in targeting cells, total RNA isolated 60 minutes post-induction was subjected to RNA sequencing (RNA-seq) and normalized 5’-end counts at each nucleotide position were compared for targeting and nontargeting samples. Overrepresented 5’-end counts were considered potential cleavage sites. Because collateral RNA degradation might cause global changes in *E. coli* transcriptome, we analyzed transcription start sites (TSSs) to rule out the possibility that overrepresented 5’-end counts detected in targeting samples were related to the start sites of transcription activated by Cas13 RNA targeting rather than cleavage sites. The top-1000 TSSs predicted in targeting samples were compared with the top-1000 predicted RNA cleavage sites. Only five sites were shared between those two sets, suggesting that the emergence of new TSSs made little if any contribution to the observed 5’ ends enrichment between targeting and nontargeting samples, and therefore validating the approach we used for the cleavage site identification (see Supplementary Information for TSS analysis).

Numerous cleavages in RNAs isolated from targeting cells were identified and mapped on *E. coli* and plasmid transcripts. The results showed that many mRNAs were prominently cut between the 2nd and 3rd nucleotides of codons located in the beginning of the coding regions (Fig. 2a and Extended Data Fig. 3a). As a representative result, Fig. 2a shows specific cleavage of plasmid-encoded *bla* transcript and genomic *rpmH* transcript at the fourth and second codons, respectively. The observed 3-nt periodicity of mRNA cleavage sites distribution along the coding regions suggested that cleavage could be coupled to the elongation of translation (Fig. 2a).

**Fig. 2:**
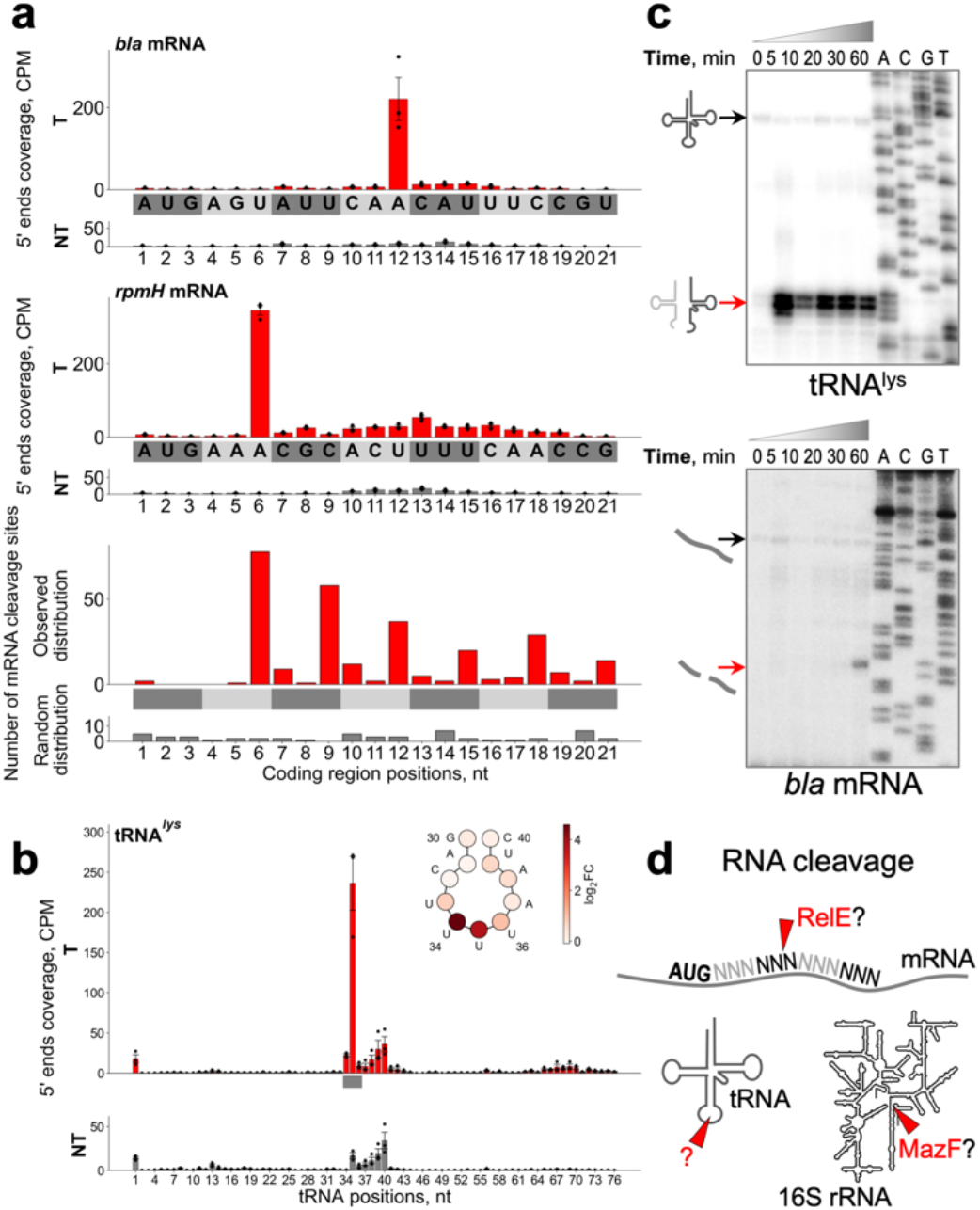
Cas13a-mediated RNA cleavage in targeting cells. **a**, CPM (counts per million) normalized coverage of 5’ ends of transcripts mapped to coding regions of *bla* (top panel) and *rpmH* (middle panel) genes in targeting (T, red bars) and nontargeting (NT, grey bars) samples of *E. coli* cells. The horizontal axes depict nucleotide positions of the coding regions. Red and grey bars correspond to mean (± s. e. m.) CPM normalized counts of mapped 5’ ends from three biological replicates. The values for individual samples are shown by dots. The bottom panel depicts the distribution of top-1000 of RNA cleavage sites (sorted by adjusted p-values in ascending order) across the positions of the coding regions fetched from the annotations. For convenient visualization, separate codons are colored in grey/dark-grey colors. **b**, CPM normalized coverage of 5’ ends of transcripts mapped onto *lysT* gene in targeting (T, red bars) and nontargeting (NT, grey bars) samples of *E. coli* cells (mean ± s. e. m. from three biological replicates). The horizontal axis depicts the nucleotide position of tRNA. The region corresponding to anticodon is shown by a grey bar. At the top right part of the panel, the scheme of *lysT* tRNA anticodon loop is depicted; circles correspond to tRNA nucleotides. Each circle is colored according to color scheme where the intensity of color corresponds to values of log_2_FC between mean 5’ ends CPMs in targeting and nontargeting samples. **c**, Kinetics of cleavage product accumulation for tRNA^lys^ and *bla* mRNA revealed by primer extension assay. Black arrows depict the position of full-size RNA, red arrows depict the position of cleavage product. The gel images are representative of three independent experiments. **d**, Schematic representation of mRNA, 16S rRNA, and tRNA cleavage in targeting cells. Cleavages in mRNA between the 2^nd^ and 3^rd^ nucleotides of a codon and in 16S rRNA removing the anti-Shine-Dalgarno sequence resemble the activity of RelE and MazF toxin RNases, respectively. For gel source data, see Supplementary Fig. 1.

Another prominent cleavage site was observed in 16S rRNA, where the cleavage removed a 3’-end fragment containing the anti-Shine-Dalgarno sequence (Extended Data Fig. 3b). Finally, massive cleavage of tRNAs in the anticodon loops was detected in targeting cells. The extent of cleavage notably varied for different tRNAs, with some, including lysine and glutamine tRNAs, being cut extensively, whereas others remained largely intact (Fig. 2b and Extended Data Fig. 4). Overall, the cleavage of mRNAs, 16S rRNA, and tRNAs appears to be responsible for the inhibition of protein synthesis induced by Cas13a RNA-targeting.

A common feature of type VI effectors is the presence of two HEPN domains required for RNase activity^8^. Each HEPN domain contains a catalytic dyad of conserved arginine and histidine residues^3,13^. Substitutions of any of these residues abolish the Cas13 RNase activity *in vitro* and protection against RNA phages *in vivo*^3^. Cultures of targeting cells expressing Cas13a with single amino acid substitutions in the HEPN catalytic residues grew at the same rate as nontargeting control cultures (Extended Data Fig. 5a-c). As expected, HEPN mutations had no effect on crRNA processing, which is known to depend on a distinct RNase active site of Cas13a^11,14^ (Extended Data Fig. 5d). In contrast, primer extension analysis showed that inactivation of the HEPN RNase abolished target-activated tRNA and mRNA cleavage (Extended Data Fig. 5e). Thus, Cas13a HEPN RNase is responsible, directly or indirectly, for the RNA cleavage observed in targeting cells.

The kinetics of accumulation of tRNA and mRNA cleavage products differed substantially (Fig. 2c). While tRNA cleavage was readily detectable as early as 5 minutes post-induction, consistent with rapid protein synthesis inhibition (Extended Data Fig. 2), mRNA cleavage products accumulated much later and became detectable ~60 minutes post-induction. Such different cleavage kinetics suggested that mRNA cleavage could be a consequence of tRNA cleavage and/or translation inhibition followed by activation of cellular RNases.

## tRNA cleavage is unaffected in targeting cells lacking RNase toxins

The cleavage of mRNA and 16S rRNA in targeting cells resembles that induced by RNase toxins, such as RelE and MazF^15–18^ (Fig. 2d). To elucidate potential connections between Cas13a and toxin RNases, we used Δ10, an *E. coli* strain that lacks 10 endoribonuclease-encoding type II toxin-antitoxin systems^19^. Wild-type and Δ10 cells were transformed with a plasmid for inducible expression of RFP and the targeting or nontargeting variant of the *L. shahii* type VI-A CRISPR-Cas system plasmid. Suppression of the Δ10 culture growth was observed upon induction of RFP mRNA targeting (Fig. 3a). RNA cleavage was next monitored by primer extension in wild-type and Δ10 targeting cultures. No mRNA cleavage was detected in RNA prepared from Δ10 targeting culture, suggesting that some of the deleted toxins were responsible for mRNA cleavage observed upon Cas13a RNA-targeting in wild-type cells (Fig. 3b). In contrast, tRNAs were efficiently cleaved in the Δ10 samples (Fig. 3c). In wild type cells, cleavage at the same RNA sites was observed in cells expressing crRNAs that targeted three different positions of the RFP transcript (Fig. 3b, c and Supplementary Table 1), confirming that it was target recognition as such that triggered mRNA and tRNA degradation. RNA purified from the Δ10 samples was also analyzed by RNA-seq. Massive tRNA cleavage with a pattern similar to that observed in RNA from wild-type cells was observed (Fig. 3d, e and Extended Data Fig. 4). In both cases, tRNAs were cut mainly at the first nucleotides of the anticodons (Fig. 3e and Extended Data Fig. 6). The consensus of the top-1000 cleavage sites detected in Δ10 cells (sorted by adjusted p-values) demonstrated a preference for uridine residues, in contrast to the adenine-rich consensus of the top-1000 cleavage sites from the wild-type cells (Fig. 3f). Previous analysis of LshCas13a collateral damage specificity *in vitro* also detected a marked preference for uridine residues at the cleavage site^3^. Thus, the A-rich cleavage site consensus in targeting wild-type cells most likely reflected the preferences of the RNases encoded by toxin-antitoxin modules.

**Fig. 3:**
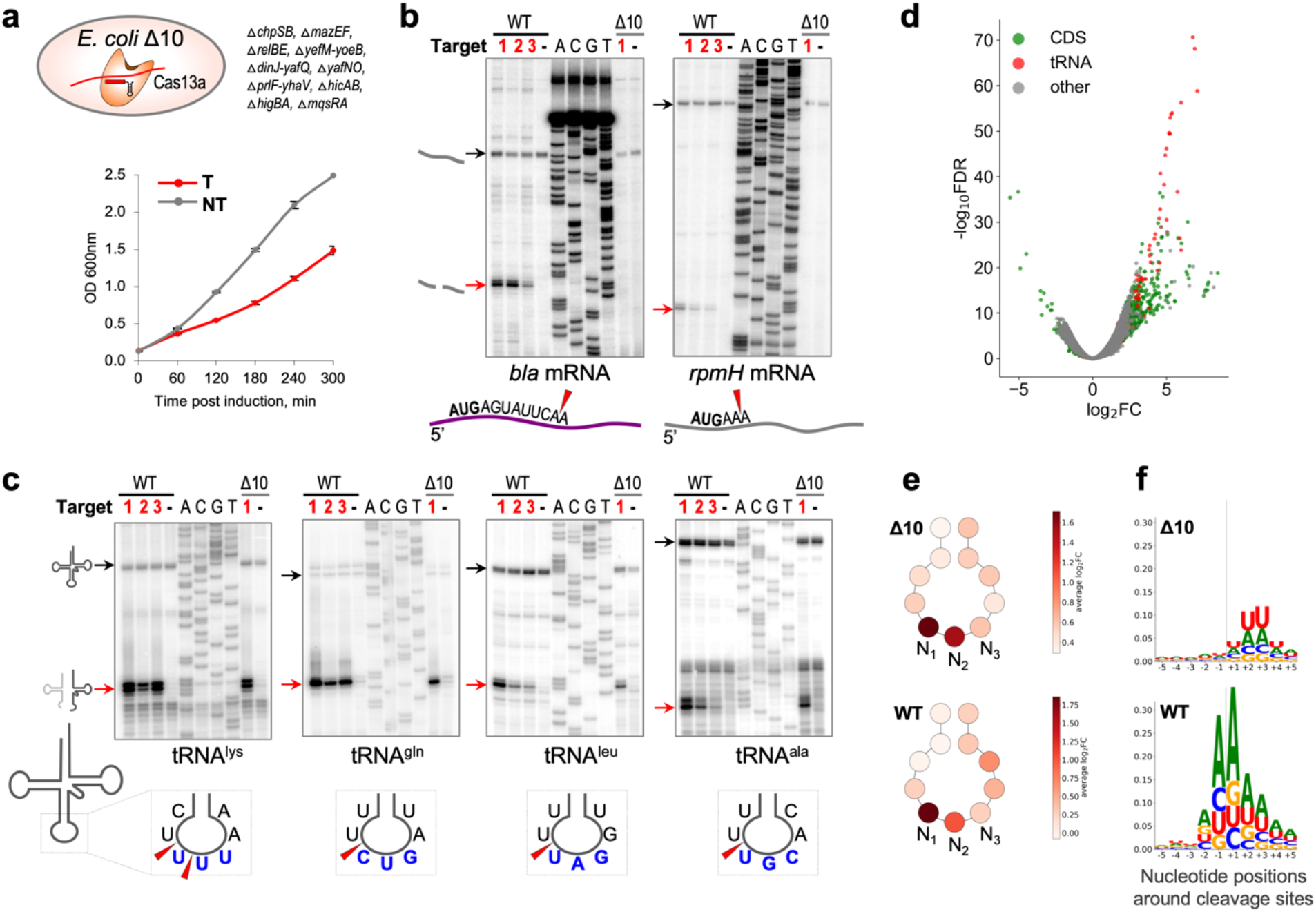
tRNAs but not mRNAs are cleaved in targeting cells lacking 10 toxin RNases. **a**, RFP mRNA targeting by LshCas13a expressed in *E. coli* Δ10 results in growth suppression (T, red); no such effect is observed in nontargeting (NT, grey) control, same as shown in wild type *E. coli* (**Fig. 1c**). OD_600_ measurements taken after RFP induction represent a mean ± s. e. m. of three biological replicates at each time point. **b**, Primer extension assay revealed mRNA cleavage in the presence of toxin RNases, but not in *E. coli* Δ10. **c**, tRNA cleavages are intact in the cells lacking toxin RNases. Black arrows depict the position of full-size RNA, red arrows depict the position of cleavage product. The gel images are representative of three independent experiments. **d**, Volcano plot depicting the results of the comparison of 5’ transcript ends counts per nucleotide position of each strand between targeting and nontargeting samples of Δ10 *E. coli* strain using likelihood ratio test. Each dot corresponds to analyzed nucleotide position. The horizontal axis depicts log_2_FC value between targeting and nontargeting samples; the vertical axis depicts adjusted p-value. Dot colors correspond to genomic features where the analyzed nucleotide position is located. **e**, Schematic depiction of tRNA anticodon loop; circles correspond to tRNA nucleotides. Each circle is colored according to color scheme where the intensity of color corresponds to the average value of log_2_FC between average 5’ ends CPMs in targeting and nontargeting samples across all tRNA genes detected in *E. coli* genome. N1, N2, N3 nucleotides correspond to tRNA anticodon. **f**, Weblogo plots built from 10-nt sequences surrounding top-1000 of RNA cleavage sites (sorted by adjusted p-values in ascending order). The position of cleavage is depicted by dash lines. In **e** and **f**, the top (WT) and the bottom (Δ10) panels correspond to the experiments with wild type and Δ10 *E. coli* strains, respectively. For gel source data, see Supplementary Fig. 1.

## Target-activated LshCas13a cleaves tRNAs at the anticodon loop *in vitro*

Because *E. coli* K12 used throughout this work as the host strain does not encode known anticodon nucleases, we hypothesized that tRNAs were cleaved directly by Cas13a activated by the interaction with its RNA target. To test this hypothesis, we performed an *in vitro* cleavage assay followed by primer extension analysis to characterize the cleavage products (Fig. 4a). Recombinant LshCas13a loaded with crRNA was combined with the corresponding target RNA and supplemented with bulk *E. coli* tRNA; nontargeting control reactions were performed in the absence of the target RNA. Cleavage of tRNAs at the same anticodon loop sites as observed *in vivo* was detected in the presence of crRNA, and a mutation of a HEPN catalytic residue abolished this cleavage (Fig. 4b). We next asked whether LshCas13a could specifically cleave tRNA *in vitro* in the presence of other cellular RNAs. We repeated the cleavage assay using total *E. coli* RNA as the cleavage substrate and detected the same tRNA cleavage products (Extended Data Fig. 7a). Thus, LshCas13a is an anticodon tRNase activated by target recognition.

**Fig. 4:**
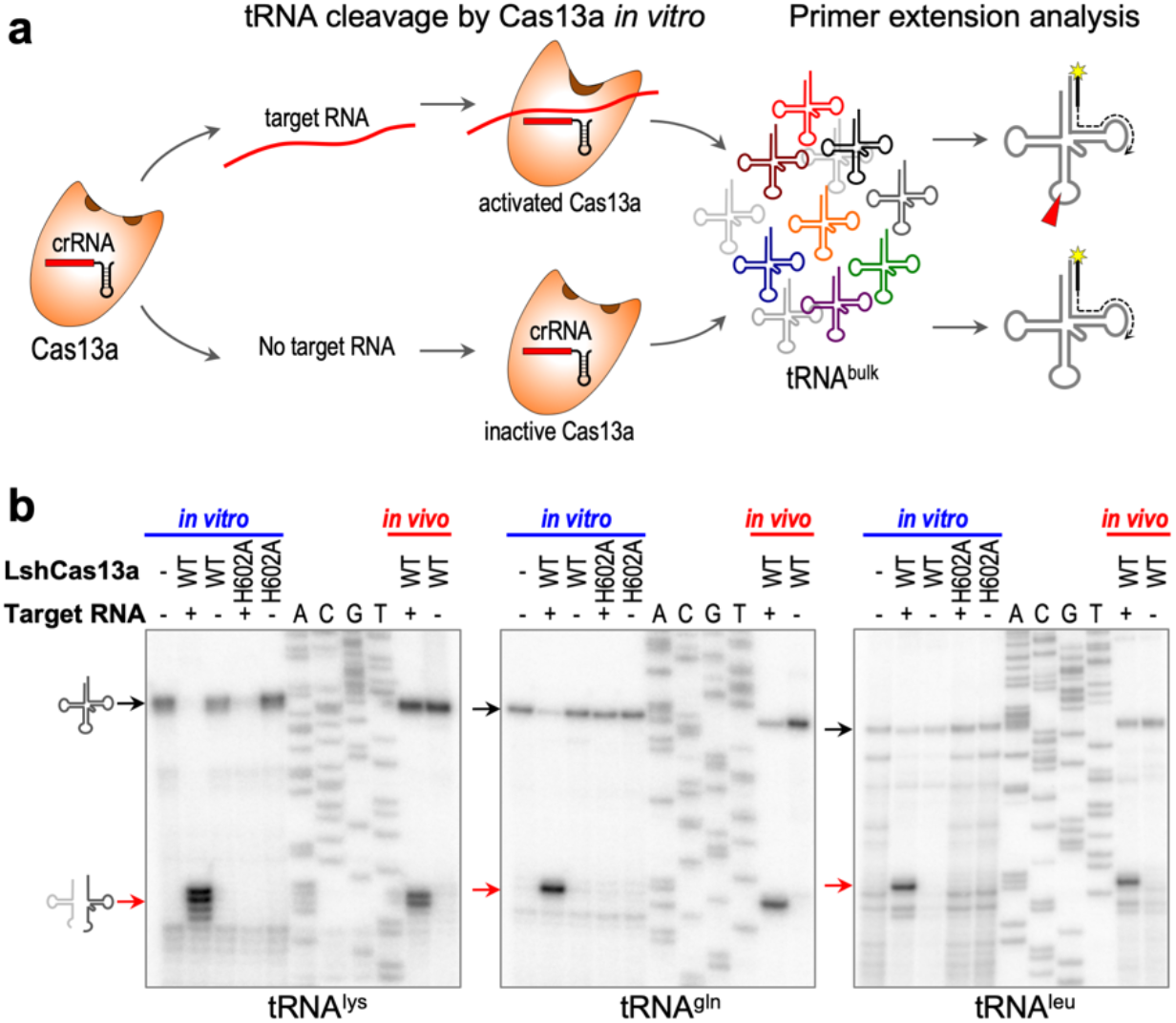
Target-activated Cas13a cuts tRNA at the anticodon loop *in vitro*. **a**, Schematic presentation of *in vitro* cleavage assay: target-activated Cas13a or inactive control are supplemented with bulk tRNA for cleavage. Cleavage products analyzed by primer extension assay using radio-labelled primer complementary to 3’-end of specific tRNA. **b**, Cleavage products revealed by primer extension following *in vitro* cleavage by target-activated LshCas13a match those observed *in vivo*. As expected, the H602A mutation of the catalytic residue in the HEPN1 domain abolishes LshCas13a cleavage activity. Black arrows show full-size tRNAs, red arrows indicate cleavage products. The gel images are representative of three independent experiments. For gel source data, see Supplementary Fig. 1.

Bacterial tRNAs are heavily modified, especially at the anticodon loops nucleotides^20^. To determine whether these modifications were essential for the Cas13a cleavage, unmodified synthetic tRNA^lys^ was tested in the *in vitro* cleavage assay. Unmodified substrate was cut by target-activated LshCas13a at the same position as natural modified tRNA (Extended Data Fig. 7b), suggesting that the specific structure of tRNAs and the anticodon loop sequence rather than base modification determine the specificity of the LshCas13a cleavage.

Total RNA was used as a substrate for LshCas13a *in vitro* cleavage followed by RNA-seq to test whether target-activated Cas13a cleaves RNAs other than tRNAs. Similar to the *in vivo* results, we detected massive tRNA cleavage at anticodon loops (Extended Data Figs. 8a, b, 9) as well as some cleavage of mRNAs and rRNAs (Extended Data Figs. 8a). However, in cells, the abundance of these non-tRNA cleavage products was negligible compared to tRNA cleavage products (Fig. 3d). A closer examination of the minor non-tRNA cleavage products showed that LshCas13a cut them at uridine residues located in small loops, apparently mimicking the anticodon stem-loop structure of tRNA (Extended Data Fig. 10).

## Discussion

Our results lead to the following view of type VI-A immunity (Fig. 5). If phage-infected cells carrying a VI-A locus express a crRNA(s) targeting a phage transcript, target recognition activates the HEPN domains of Cas13a enabling collateral RNA cleavage. Target-activated Cas13a cleaves a subset of tRNAs at their anticodon loops. tRNAs containing U-rich anticodons are preferred substrates for cleavage reflecting the specificity of the activated Cas13a tRNase (Extended Data Fig. 6). Massive cleavage of tRNAs leads to ribosome stalling, which activates type II toxin RNases cleaving mRNA and rRNA. Degradation of mRNAs and rRNAs by these toxins further compromises translation. Although there is an obvious link between RNA-targeting by Cas13a and toxin RNase activities, direct cleavage of tRNAs by target-activated Cas13a is sufficient to arrest protein synthesis and suppress cell growth. Thus, tRNA cleavage is probably the primary mechanism through which type VI CRISPR-Cas systems induce dormancy and limit the spread of phage infection through the population, whereas the activation of TAs is a secondary, back-up mechanism.

**Fig. 5:**
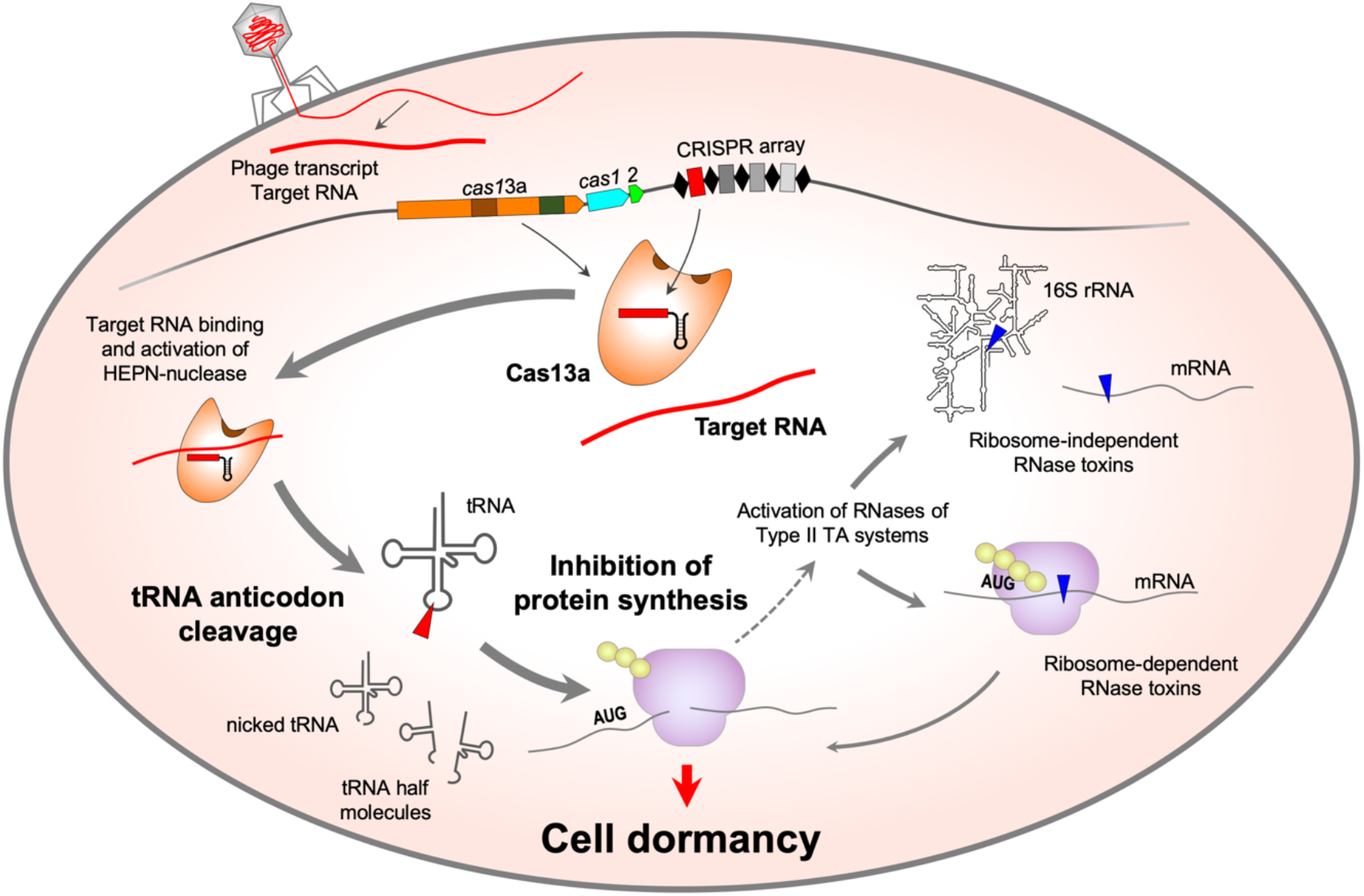
Mechanism of CRISPR-Cas immunity in type VI-A system of *Leptotrichia shahii.*

The type VI CRISPR systems are thought to have evolved from an abortive infection (Abi) system containing a HEPN RNase^6,7,13^. Our present results imply that the ancestral Abi nuclease functioned by abrogating translation via tRNA cleavage. The combination of such a nuclease with CRISPR array and adaptation module endowed the system with increased specificity and hence reduced the fitness cost incurred by the toxic Abi module.

Depletion of specific tRNAs leads to ribosome stalling at ‘hungry’ codons, which can trigger frameshift or ribosome bypass and cause synthesis of aberrant polypeptides. Bacterial anticodon nucleases, such as PrrC and colicins D and E5, apparently cause this outcome because they completely deplete the pool of specific tRNAs^21–23^. In contrast, our results demonstrate that the Cas13a-mediated tRNA cleavage does not result in complete depletion of full-size tRNAs, at least in the surrogate system used here (Fig. 3c, Extended Data Fig. 4) although the effect sufficed for translation arrest. The tRNA fragments produced by Cas13a cleavage might act as signaling or inhibitory molecules.

Bacteriophage RNA ligases that rejoin tRNA halves produced by anticodon nucleases, rendering phages resistant to this defense mechanism^24,25^, as well as bacterial tRNA ligases whose expression is activated by tRNA fragments^26^ are known. Such RNA ligases could function as anti-CRISPR mechanisms against type VI, in which case the activation of RNase toxins would become the primary defense line. On the practical side, determination of preferred collateral cleavage substrates for target-activated Cas13a effectors could help improving the sensitivity of the powerful SHERLOCK method of nucleic acids detection^27^.

## METHODS

### *E. coli* strains and plasmids

*E. coli* C3000 (wild-type) cells were transformed with two plasmids: CRISPR Cas13 plasmid containing the *Leptotrichia shahii* (Lsh) Cas13a locus and RFP plasmid for inducible expression of red fluorescent protein (RFP). The targeting cells contained the CRISPR Cas13a plasmid encoding crRNA spacer targeting the RFP mRNA. The cells contained the CRISPR Cas13a plasmid encoding crRNAs with no matching sequences in *E. coli* genome and the plasmids were used as the nontargeting control. CRISPR Cas13 plasmids (pC002, pC003, pC008, and pC003_RFP1) described previously^3^ were gifts from the Feng Zhang lab. Additional CRISPR Cas13a plasmids were constructed using Golden Gate cloning on the base of pC003 plasmid containing BsaI sites for spacer cloning (Supplementary Table 1). All CRISPR Cas13 plasmids have a pACYC184 backbone carrying a chloramphenicol resistance gene. RFP plasmid (pC008) is pBR322 derivative carrying *bla* gene for ampicillin resistance and *rfp* gene which expression is under control of tetR-promoter induced by anhydrotetracycline. *E. coli* Δ10 strain contained genomic deletions of 10 type II toxin-antitoxin systems (Δ*chpSB,* Δ*mazEF,* Δ*relBE,* Δ*yefM-yoeB,* Δ*dinJ-yafQ,* Δ*yafNO,* Δ*prlF-yhaV,* Δ*hicAB,* Δ*higBA,* Δ*mqsRA*) described previously^19^ was a gift from the Van Melderen lab. *E. coli* Δ10 strain was transformed with two plasmids: CRISPR Cas13 plasmid containing the LshCas13a locus encoding Cas13a and crRNA spacer targeting the RFP mRNA, and RFP plasmid for inducible expression of RFP to construct targeting Δ10 cells. In nontargeting Δ10 cells, CRISPR Cas13 plasmid encodes crRNA with no matching sequences in *E. coli* genome and the plasmids.

### Cell growth assay

To investigate the effect of the RFP mRNA targeting on the cell growth rate, individual colonies of transformed targeting and nontargeting wild-type or Δ10 *E. coli* cells were grown overnight in LB supplemented with 25 μg/ml chloramphenicol and 100 μg/ml ampicillin. Cell cultures were diluted 1:100 in fresh LB containing chloramphenicol and ampicillin and grown for 1 hour at 37°C with continuous shaking. After 1 hour, RFP expression was induced with 500 ng/ml anhydrotetracycline and OD_600_ measurements were taken every 30 minutes. All growth experiments were performed at least three times.

### Time-lapse microscopy

For microscopy, *E. coli* cells supplemented with CRISPR Cas13 plasmid and RFP plasmid were grown under the same conditions as described above – overnight cultures were diluted 1:100 and grown for 1 hour at 37°C. Aliquots of the cell cultures mixed with 100 nM YOYO-1 dye were dropped on an LB-1.5% agarose block supplemented with 25 μg/ml chloramphenicol and 100 μg/ml ampicillin, and 500 ng/ml anhydrotetracycline for induction of RFP expression. Two agarose blocks containing targeting and nontargeting cells were placed into one microscope chamber and cell growth of two cultures was simultaneous monitored under induced conditions. The experiment was done in triplicates. The Nikon Ti-E inverted microscope was equipped with Andor’s Zyla 4.2 sCMOS camera, Semrock filter Set YFP-2427B for green fluorescence detection and custom-made incubation system to maintain cells at 37°C. Image analysis was done using ImageJ software^28^. For each of the three replicates, the fate of at least 100 cells were monitored, and two cell types were determined - dividing cells that formed micro-colonies over the course of the experiment, and non-dividing cells that did not form colonies nor had any visible changes in cell morphology. Both cell types were represented as a percentage of the total number of cells counted for each replicate. To distinguish between live and dead cells, green-fluorescent membrane-impermeant YOYO-1 dye was used. The dye cannot penetrate live cells but can penetrate dead cells to stain the DNA, making dead cells fluoresce green. Number of dead cells were counted in targeting and nontargeting cultures and at least 100 cells were analyzed individually for three replicates.

### Metabolic labelling and autoradiography

Targeting and nontargeting *E. coli* C3000 cells were grown in M9 minimal media containing 18 amino acids without methionine and cysteine till 0.1 OD_600_ at 37°C with continuous shaking. RFP expression was then induced with 500 ng/ml anhydrotetracycline. 500 μl-aliquots were taken at time 0 and then at 10, 30, 60, and 120 min post-induction and mixed with 25 μl of a solution of thymidine (10 μg/ml), or uridine (50 μg/ml), or a mixture of methionine and cysteine (10 μg/ml each) containing 1 μCi of radioactive [methyl-^3^H]-thymidine (6.7 Ci/mmol), or [5,6-3H]-uridine (35-50 Ci/mmol), or L-[^35^S]-methionine (1135 Ci/mmol), (Perkin-Elmer, USA), respectively. Pulse-labelling was carried out at 37°C for 2 min with continuous shaking at 300 rpm, followed by the addition of 100 μl cold 40% TCA to stop the reaction. Samples were filtered through glass microfiber filters (GE Healthcare Whatman), and the filters were washed twice with 1 ml cold 10% TCA and 1 ml cold 100% ethanol each. The dried filters were placed in 4 ml of scintillation liquid, and the radioactivity was measured in a liquid scintillation counter (LS60001C, Beckman Coulter, USA). Radioactivity counts corresponding to incorporated thymidine, uridine, and methionine were normalized to OD450 at each time point. For the analysis of RFP expression relative to overall protein synthesis, the targeting and nontargeting cells were grown as described above, except that 500 μl-aliquots for [^35^S]-methionine pulse-labeling reactions were taken before (time 0) and after RFP induction at 2, 5, and 10 min. The samples were analyzed by MOPS/SDS 10% polyacrylamide gel electrophoresis (PAGE) followed by Coomassie staining and quantification by Phosphorimager.

### RNA isolation

Total RNA was isolated from *E. coli* cells growing in LB media and harvested at 60 min post-induction of RFP expression. Cell lysis was done using Max Bacterial Enhancement Reagent (Invitrogen) for 4 min and then with TRIzol reagent (Invitrogen) for 5 min. RNA was extracted by chloroform and precipitated with isopropanol. RNA pellets were washed with 70% ethanol and then dissolved in nuclease free water and then treated with Turbo DNA-free kit (Invitrogen) to remove DNA contamination.

### Primer extension

For primer extension, 3.5 μg total RNA was reverse transcribed with the SuperScript IV First-Strand Synthesis System (Invitrogen) according to the manufacturer’s protocol. DNA oligonucleotides (Supplementary Table 2) were radiolabeled at the 5’ end by T4 Polynucleotide kinase (PNK) (New England Biolabs) treatment and [γ-^32^P] (PerkinElmer) for 60 min at 37°C followed by purification on a Micro-Bio Spin P-6 Gel Column (Bio-Rad). 1 pmol of 5’-end-radiolabeled primer was used for each extension reaction. In parallel, sequencing reactions were performed on amplified PCR fragments of genomic or plasmid loci with the corresponding radiolabeled primers using the Thermo Sequenase Cycle Sequencing Kit according to manufacturer’s instructions. Reaction products were resolved by 10% denaturing PAGE and visualized using a Phosphorimager.

### HEPN mutagenesis

Alanine mutations were created in each of four catalytic residues of LshCas13a HEPN domains (R597A, H602A, R1278A and H1283A) in plasmid containing LshCas13a locus and a CRISPR array carrying a spacer targeting RFP mRNA (pC003_RFP1) using QuikChange Site-Directed Mutagenesis kit (Agilent) and the mutagenic primers containing the desired mutations (Supplementary Table 2) according to the manufacturer’s protocol. Cell growth experiments were done as described above. Primer extension analysis to determine cleavage products in *bla* mRNA, *rpmH* mRNA and tRNAs was also done as described above. In all HEPN mutant experiments, wild-type targeting and nontargeting variants were used as controls.

### Northern blot hybridization

The procedure was mostly performed as described before^29^. 10 μg of total RNA from *E. coli* cells transformed with CRISPR Cas13 plasmid expressing HEPN mutant LshCas13a proteins with mutations R597A, H602A, R1278A, H1283A, were resolved by 10% denaturing PAGE in Mini Protean 3 Cell (Bio-Rad). Separated RNA was then transferred to a nylon membrane (Hybond-XL, GE Healthcare) in pre-chilled 0.5xTBE buffer for 90 min at 50 V using Mini Trans-Blot electrophoretic transfer cell (Bio-Rad). Transferred RNA was UV-crosslinked to membrane and hybridized with [γ- ^32^P] 5’-end labeled RFP-spacer specific probe (RFP_crRNA_probe, Supplementary Table 2) in ExpressHyb solution (Clontech Laboratories, Inc) according to manufacturer’s protocol. Hybridization was performed in Isotemp rotisserie oven (Fisher Scientific) for 2 hours at 37°C. Hybridized RNA bands were visualized by phosphorimaging.

### LshCas13a protein purification

Lsh *cas13a* gene was cloned into NdeI and BamHI sites in pET28 plasmid for protein purification. *E. coli* BL21(DE3) cells were transformed with pET28_LshCas13a for protein expression and isolation. LshCas13a protein was purified from the cells grown in 400 ml LB supplemented with 50 μg/ml kanamycin and 0.5 mM IPTG, as described previously^30^. Freshly transformed cells were grown at 37°C to OD_600_ 0.6–0.9, then induced with 0.5 mM IPTG and grown for additional 6–8 hours at room temperature before cell harvesting. The cell pellets resuspended in buffer A (20 mM Tris-HCl pH, 8.0, 500 mM NaCl, 4 mM imidazole pH 8.0, 5% (v/v) glycerol, 0.2 μg/ml phenylmethylsulfonyl fluoride (PMSF)) containing protease inhibitor cocktail Roche cOmplete, EDTA-free (Sigma) were lysed by sonication. Lysates were cleared by centrifugation at 15,000 g for 60 min and filtered through 0.22 micron filter (Millipore) and applied to a 1-ml chelating Hi-Trap Sepharose column (GE Healthcare) equilibrated with buffer A. Proteins were purified first by washing the column using Buffer A containing 25 mM imidazole and then eluting with buffer A containing 200 mM imidazole. Pooled protein fractions were diluted 10 times with TGED buffer (20 mM Tris-HCl, pH 8.0, 5% (v/v) glycerol, 1 mM EDTA, 2 mM β-mercaptoethanol) and applied to a 1-ml Hi-Trap Heparin column (GE Healthcare) equilibrated with TGED. The column was washed with TGED containing 500 mM NaCl, and proteins were eluted with TGED containing 1 M NaCl. Pooled protein fractions were concentrated using Microsep centrifugal devices 30K (Pall Corp), dialyzed against buffer B (20 mM Tris-HCl, pH 8.0, 200 mM NaCl, 50% (v/v) glycerol, 0.5 mM EDTA, 2 mM β-mercaptoethanol) and then stored at −80°C. HEPN H602A encoding mutation was introduced into pET28_LshCas13a plasmid using QuikChange Site-Directed Mutagenesis kit as described above and mutant LshCas13a was purified similar to the wild-type LshCas13a.

### Generation of RNA for *in vitro* cleavage assay

crRNA and target RNA were transcribed *in vitro* from PCR-generated dsDNA template using T7 RNA polymerase (New England Biolabs) according to manufacturer recommendations and was purified by electrophoresis in 10% polyacrylamide 6M urea gels. Oligonucleotide sequences are listed in Supplementary Table 2. Bulk *E. coli* tRNA was ordered from Sigma. Unmodified tRNA^lys^ was ordered synthetically (Integrated DNA Technologies).

### *In vitro* cleavage assay

*In vitro* RNA cleavage was performed with LshCas13a at 37 °C in cleavage buffer (20 mM Tris-HCl pH 8.0, 100 mM NaCl, 5 mM MgCl_2_, 1 mM DTT). LshCas13a-crRNA complex was formed by combining, in 10 μl, Cas13a and crRNA (200 nM each) and incubating at 37 °C for 20 min. Next, 100 nM of target RNA was added. Nontargeting control reactions were performed in the absence of target RNA. Immediately after adding target RNA, the reactions were supplemented with collateral RNA. 0.1 μg of *E. coli* bulk tRNA, 2 μg of total RNA or 0.05 μg of unmodified synthetic tRNA^lys^ per 10 μl reaction were used as collateral cleavage substrates. After 60 min of incubation at 37 °C, RNA from reaction mixture was extracted with chloroform and precipitated in 75% ethanol in the presence of 0.3 M NaOAc and 0.1 mg/ml glycogen. tRNA cleavage products were analyzed by primer extension as described above. For RNA sequencing analysis of cleavage products generated by LshCas13a *in vitro*, total RNA from *E. coli* cells depleted from rRNA (MICROBExpress Bacterial mRNA Enrichment Kit (Invitrogen)) was used as cleavage substrate, the cleavage reactions were performed in 40 μl in triplicates.

### RNA sequencing

The general procedure to prepare RNA for sequencing was similar to the protocol described previously^31^ with some modifications. Total RNA samples were treated with MICROBExpress Bacterial mRNA Enrichment Kit (Invitrogen) for rRNA depletion prior to library preparation. To construct libraries contained both primary transcripts carrying 5’-triphosphate (5’-P) and processed transcripts carrying 5’-monophosphate (5’-P) or 5’-hydrocyl (5’-OH), RNA samples were treated with RNA 5’ Pyrophosphohydrolase (RppH) (New England Biolabs) for 30 min at 37°C. In parallel, RNA samples without RppH treatment were used for preparation of libraries enriched in processed transcripts. Fragmentation was carried out by sonication using the Covaris protocol to obtain fragments of 200 nt. T4 PNK (New England Biolabs) treatment was performed to convert 5’-OH to 5’-P and 3’-P to 3’-OH and prepare RNA for adapter ligation during library preparation. Samples were purified using the Zymo Research Oligo Clean and Concentrator kit. Library preparation was done using the NEBNext Multiplex Small RNA Library Prep Set for Illumina according to the manufacturer’s protocol. BluePippin size selection was done using 2% agarose gel cassette (Sage Science) to select for 100 - 600 bp products. QC at each step was carried out by both Qubit and fragment analyzer. RNA-seq was performed using Illumina NextSeq High-Output kit 2 × 35 bp paired-end reads at Waksman Genomics Core Facility, Rutgers University. The similar procedure was performed for RNA library preparation and sequencing of RNA cleavage products produced by target-activated LshCas13a *in vitro*. RNA-seq *in vit*ro cleavage products was performed at Skoltech Genomics Core Facility.

### RNA sequencing data analysis

All custom scripts used in the analysis are deposed at GitHub: https://github.com/matveykolesnik/LshCas13a_RNA_cleavage

Raw RNA sequencing reads were filtered by quality with simultaneous adapters removal using trimmomatic v. 0.36^32^. The exact parameters of trimmomatic run are available in the *raw_data_processing.sh* file. Adapters content and quality of reads before and after the processing was assessed using FastQC v. 0.11.9. Processed reads were mapped onto reference sequences (RefSeq: NC_000913.3 supplemented with pC002 and pC008 plasmids for nontargeting samples and NC_000913.3 supplemented with pC003_RFP_spacer and pC008 plasmids for targeting samples), using bowtie2 v. 2.3.4.3^33^ producing corresponding SAM files. For the transcription start sites (TSS) detection, processed reads for nontargeting samples with or without RppH treatment were mapped to the NC_000913.2 sequence. To analyze RNA-seq data for *in vitro* cleavage experiments, RNA reads were mapped to the NC_000913.3 sequence. The exact parameters of bowtie2 run are available in the *read_mapping.sh* file. Next, for each nucleotide position of each strand of reference sequences, the number of 5’ ends of aligned fragments were counted, producing corresponding tables (see *return_fragment_coords_table.py* file for details). The obtained tables were joined using *merge_ends_count_tables.py* script. The differences between the numbers of mapped 5’ ends in targeting and nontargeting samples were analyzed using edgeR package v. 3.26.3^34^. The features (here, strand specific nucleotide positions) with low counts were excluded from the analysis. The trimmed mean of M-values (TMM) normalization method implemented in edgeR was applied. Next, the edgeR likelihood ratio test was performed. The obtained p-values were corrected using Benjamini-Hochberg method, and the result tables containing analyzed features with assigned log_2_FC and adjusted p-values were written to separate files (see *TCS_calling.R* file for details). The features with log_2_FC > 0 were considered as putative RNA cleavage sites. To build weblogo plots, the identified RNA cleavage sites were sorted by adjusted p-values in ascending order, and top-1000 were selected for the analysis. Ten nucleotides surrounding the selected sites were obtained from the reference sequences using Biopython toolkit^35^, and the Logomaker^36^ module was used to create weblogo plots (see *TCS_LRT_weblogo.ipynb* file for details). Secondary structures of transcripts were predicted using the RNAfold tool from the ViennaRNA package^37^ and visualized using the forgi module^38^ (see *draw_hairpin_structures.ipynb* file for details). To visualize the cleavage of different tRNAs, the sequences of tRNA genes that were included in the analysis, were extracted from annotated NC_000913.3 assembly and aligned using MAFFT v. 7.453 with --maxiterate 1000 --localpair parameters. Each position of the alignment was assigned with the corresponding -log10(adjusted p-value) depicting the statistical significance of the enrichment of 5’ ends counts in targeting samples over nontargeting samples in d10 strain. The positions corresponding to gaps or the positions excluded from the analysis were assigned with zero values. The resulting table was visualized as the heatmap where the intensity of color depicts the -log10(adjusted p-values). To validate the approach used for identification of RNA cleavage sites, we apply this method to define TSS. The differences between the numbers of mapped 5’ ends in the samples with and without RppH treatment were analyzed in the same way as it was done for the detection of RNA cleavage sites. Predicted TSSs were compared with the *E. coli* MG1655 TSSs list from RegulonDB^39^.

**Supplementary Information** is available and contains a Supplementary Discussion, Supplementary Tables 1 and 2, Supplementary Figures 1 and 2, gel source data, and Supplementary analysis.

## Data availability

Sequencing data and custom scripts used in the analysis are available at the NCBI Gene Expression Omnibus (GEO Series GenBank accession number GSE183061) and at GitHub: https://github.com/matveykolesnik/LshCas13a_RNA_cleavage

## Acknowledgements

We thank Dr. Dibyendu Kumar and Dr. Min Tu from Waksman Genomics Core Facility, Rutgers University and Maria Logacheva from Skoltech Genomics Core Facility for performing high-throughput sequencing for this project. The microscopy experiments were carried out using scientific equipment of the Center of Shared Usage “The analytical center of nano- and biotechnologies of SPbPU”. This work was supported by NIH grant GM10407 (K.S.), Saint Petersburg State University grant ID 73450983 (A.S.), and intramural funds from the Department of Cell Biology and Neuroscience at Rowan University (S.B.); RNA-seq *in vit*ro cleavage products was supported by the Skoltech Life Sciences Program grant.

## Author contributions

E.S. and K.S. conceived the study and designed experiments with input from E.V.K., K.S.M. and S.B.; M.K. performed bioinformatic analysis. I.J., L.M., N.M., A.S., A.K., K.K., S.B. and E.S. executed all experimental work. M.K., S.M., I.J., S.B., K.S.M. and E.S. analyzed the data. K.S., E.V.K. and E.S. wrote the manuscript. All authors discussed the data and revised the manuscript.

---

**Extended Data Fig. 1.**
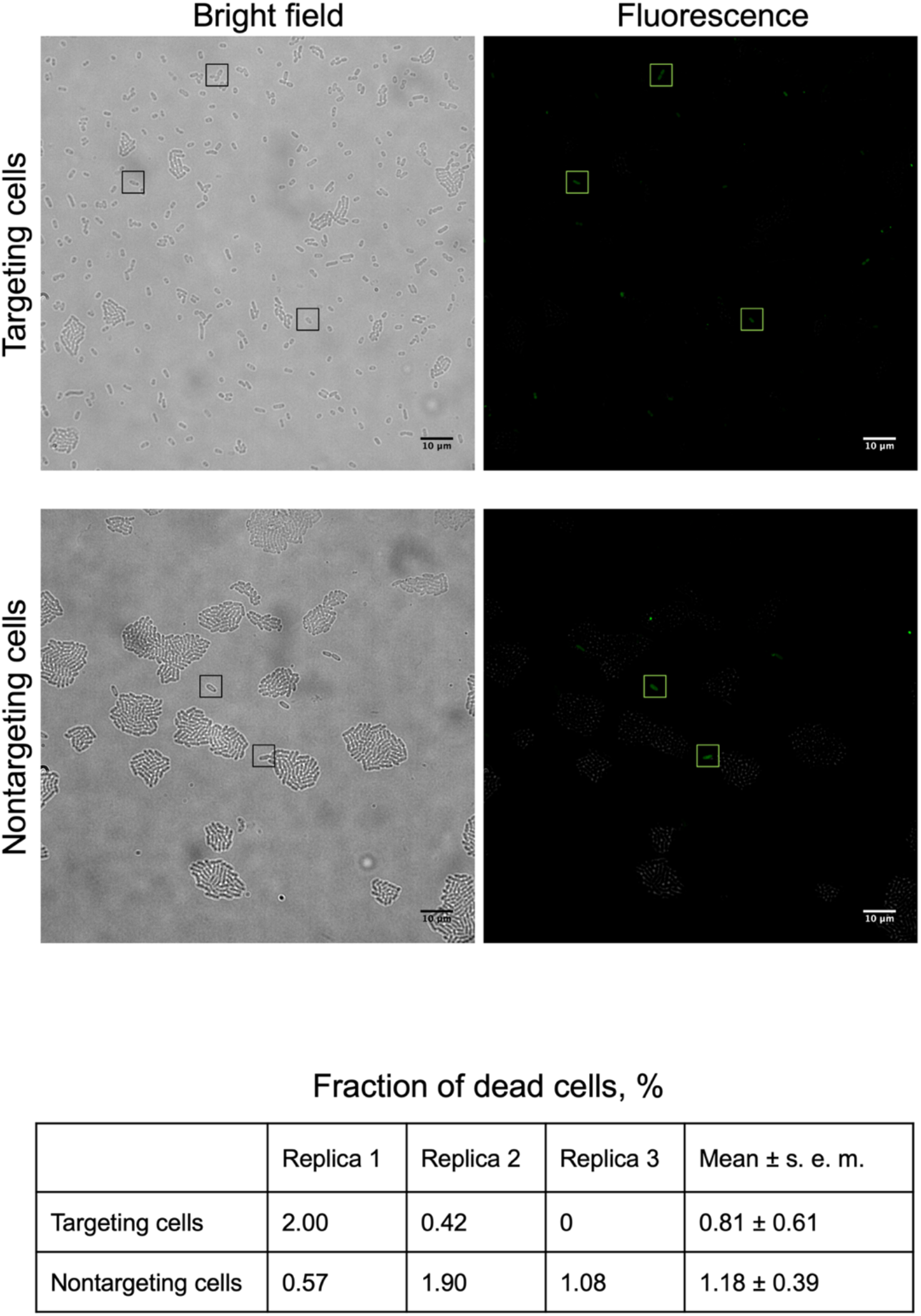
YOYO-1 staining revealed a small number of dead cells equally in targeting and nontargeting cultures. Representative bright-field and fluorescence images of targeting and nontargeting cultures stained with a membrane-impermeable dye YOYO-1 for detection of dead cells. Bright-field and fluorescence images were acquired for the same field of view at 168 minutes post-induction. Small boxes indicate dead cells fluorescing green due to YOYO-1 bound to DNA. Statistics for dead cells’ fraction was calculated from three biological replicates for targeting and nontargeting cultures. At least 100 cells were analyzed for each experiment.

**Extended Data Fig. 2.**
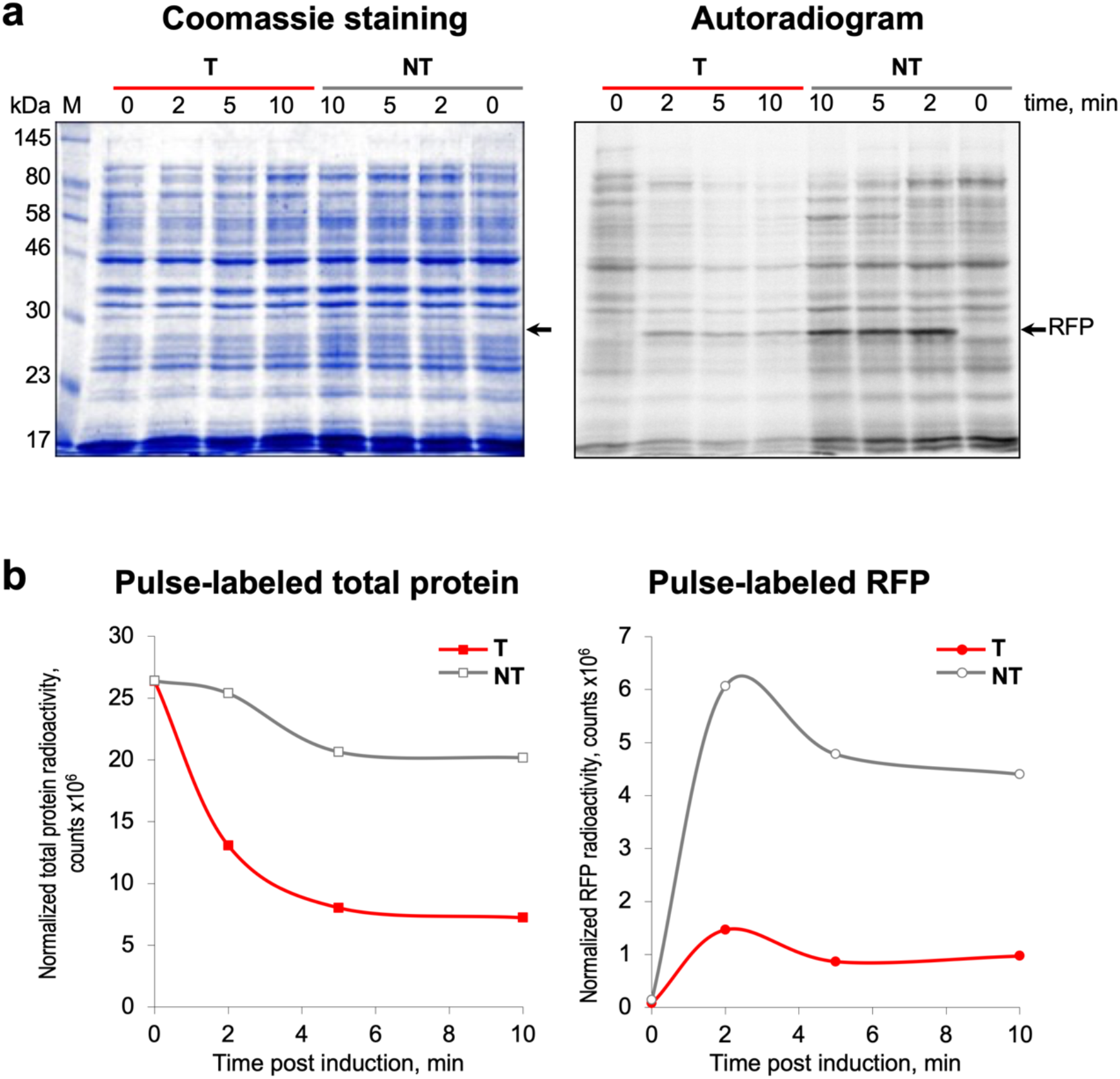
Time course of protein synthesis in targeting (T) and nontargeting (NT) cells after RFP induction. **a**, Protein samples obtained during pulse-labeling of *E. coli* cells by [^35^S]-methionine, were separated by MOPS/SDS 10% PAGE, gels were stained by Coomassie, and quantified by Phosphorimager. The position of RFP on the gel and the autoradiogram is indicated by a black arrow. **b**, Graphs plots show the ^35^S-radioactivity of pulse-labeled RFP and total cellular proteins in targeting (T, red) and nontargeting cells (NT, grey). For gel source data, see Supplementary Fig. 2.

**Extended Data Fig. 3.**
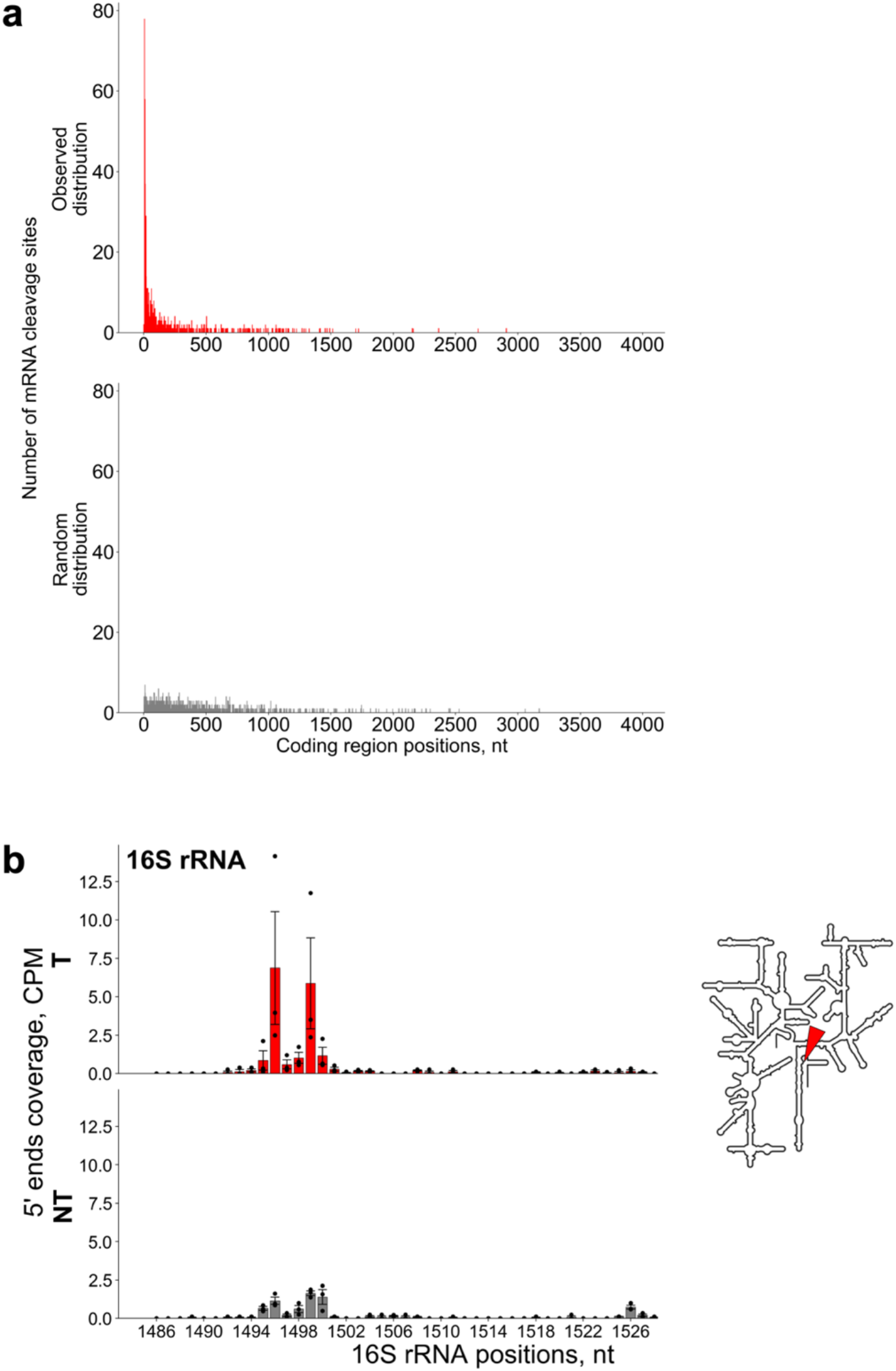
mRNA and 16SrRNA cleavages upon RFP mRNA targeting by Cas13a. **a**, The distribution of top-1000 of mRNA cleavage sites (sorted by adjusted p-values in ascending order) across the positions of the coding regions revealed efficient cleavage at the beginning of coding sequences. **b**, 16S rRNA cleavage in targeting cells. Cut of 3’-end fragment containing anti-Shine-Dalgarno sequence or cleavage at this position in rRNA precursor. CPM normalized coverage of 5’ ends of 16S rRNA in targeting (T, red bars) and nontargeting (NT, grey bars) samples of *E. coli* cells (mean ± s. e. m. from three biological replicates). The horizontal axis depicts the nucleotide position of 16S rRNA.

**Extended Data Fig. 4.**
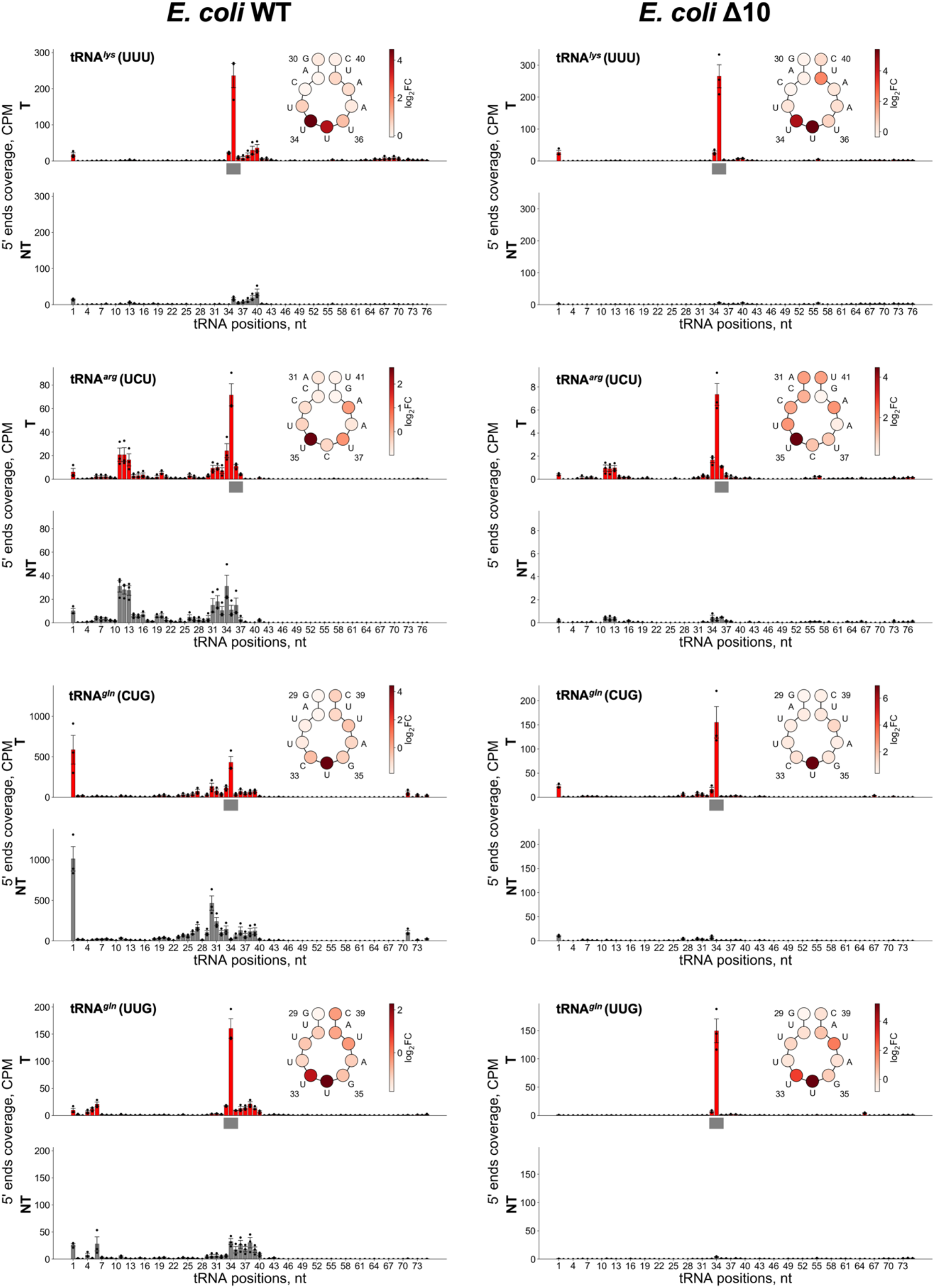
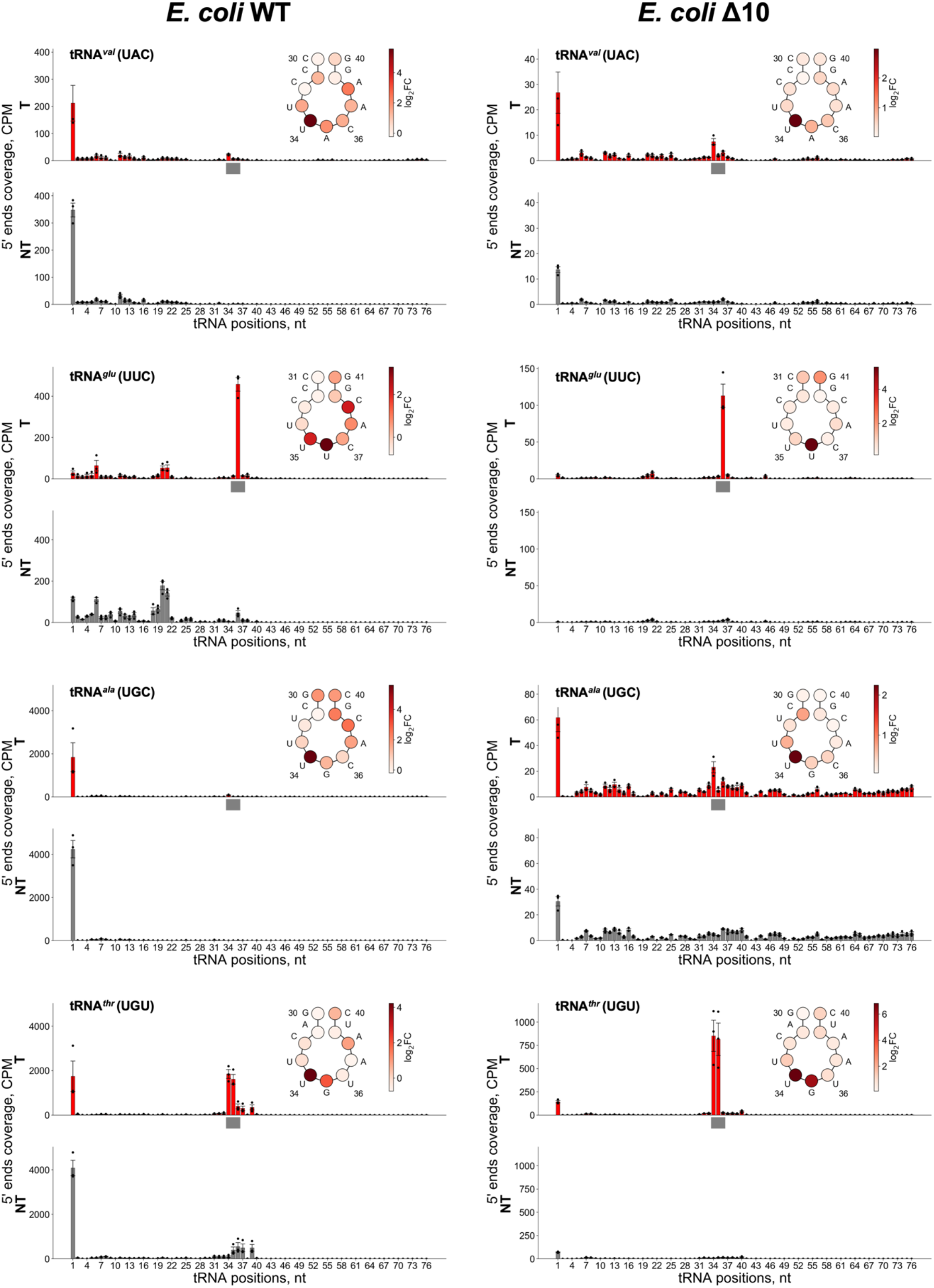
Visualization of cleavage sites in tRNAs in wild type and Δ10 *E. coli* cells. The analysis was performed in the same way as for Fig. 2b: CPM normalized coverage of 5’ ends of transcripts mapped onto tRNA gene in targeting (T, red bars) and nontargeting (NT, grey bars) samples of *E. coli* cells (mean ± s. e. m. from three biological replicates). The horizontal axis depicts the nucleotide position of tRNA. The region corresponding to the anticodon is shown by a grey bar. At the top right part of each panel the scheme of tRNA anticodon loop is depicted; circles correspond to tRNA nucleotides. Each circle is colored according to color scheme where the intensity of color corresponds to values of log_2_FC between mean 5’ ends CPMs in targeting and nontargeting samples.

**Extended Data Fig. 5.**
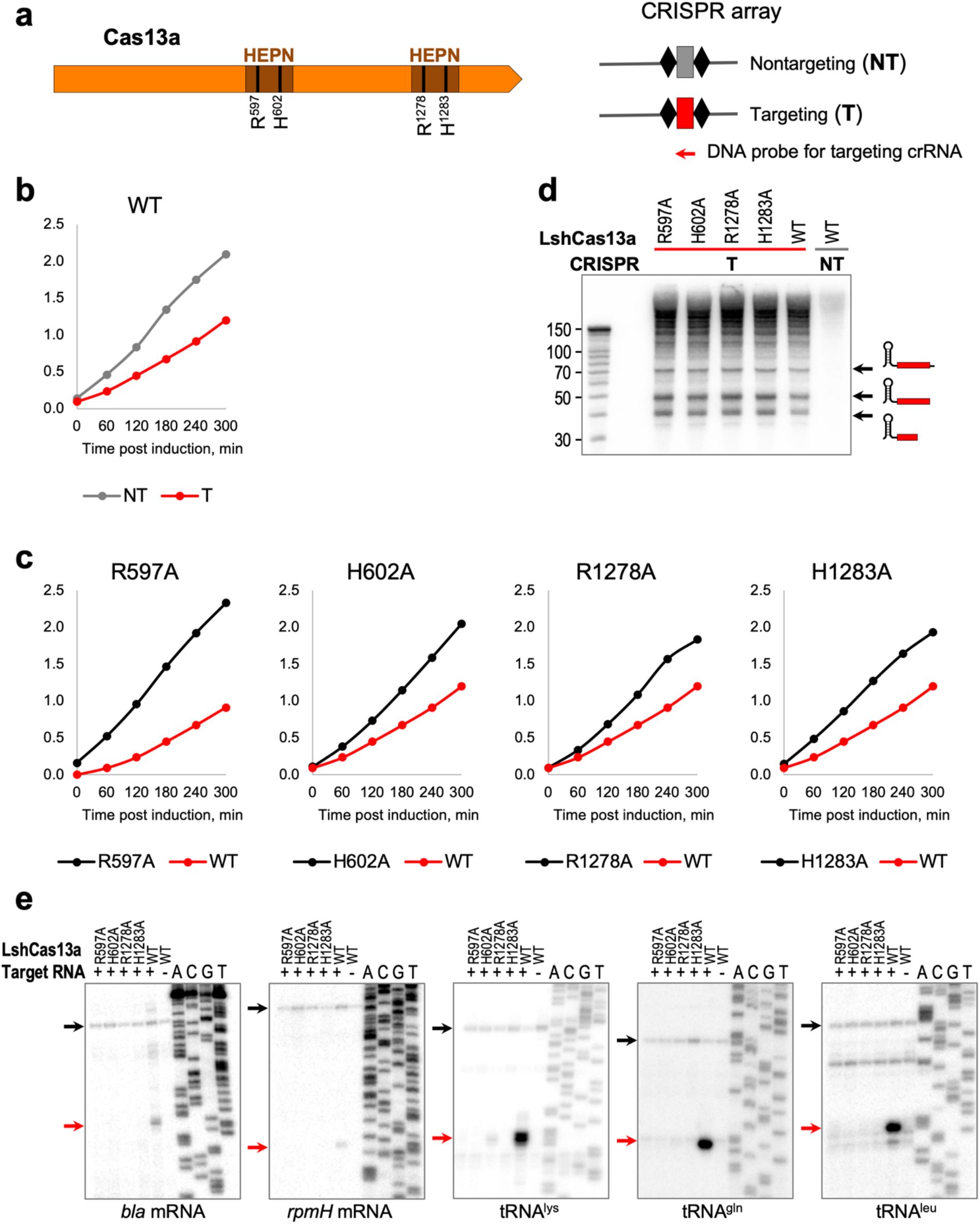
Effect of HEPN catalytic residue mutations on cell growth, crRNA processing, and RNA cleavage in cells expressing target-activated Cas13a. **a**, Schematic of the domain organization of LshCas13a protein showing catalytic residues of HEPN domains. CRISPR array structures for cells used in these experiments are shown on the right. **b**, **c**, Single substitution of any of four catalytic residues restores growth in cells expressing mutant Cas13a, crRNA and complementary target RNA, the growth rate is similar to that of nontargeting control cells expressing wild type Cas13. **d**, HEPN mutations do not affect crRNA processing. **e**, HEPN mutations abolish mRNA and tRNA cleavages. All gels are representative of three biological replicates. For gel source data, see Supplementary Fig. 2.

**Extended Data Fig. 6.**
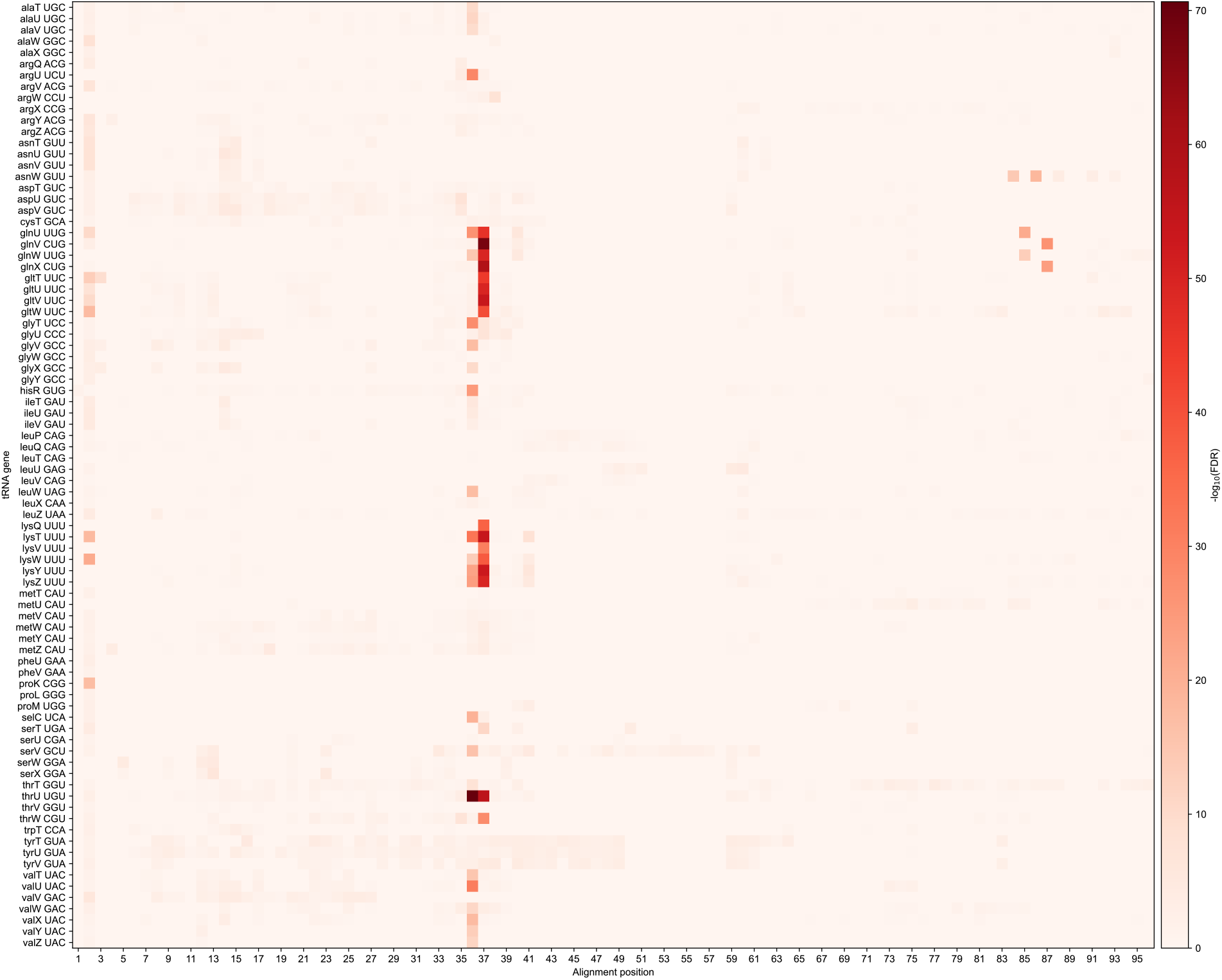
The preference of the cleavage of different tRNAs in Δ10 strain. The nucleotide sequences of tRNA genes were aligned. The horizontal axis depicts the position of the multiple sequence alignment. The vertical axis corresponds to different tRNA genes. The positions of each tRNA in the alignment were assigned with the corresponding −log10(adjusted p-value) that depicts the statistical significance of the cleavage in the given position. The positions corresponding to gaps or the positions excluded from the analysis were assigned with zero values. The resulting table was visualized as the heatmap where the intensity of color depicts the −log10(adjusted p-value).

**Extended Data Fig. 7.**
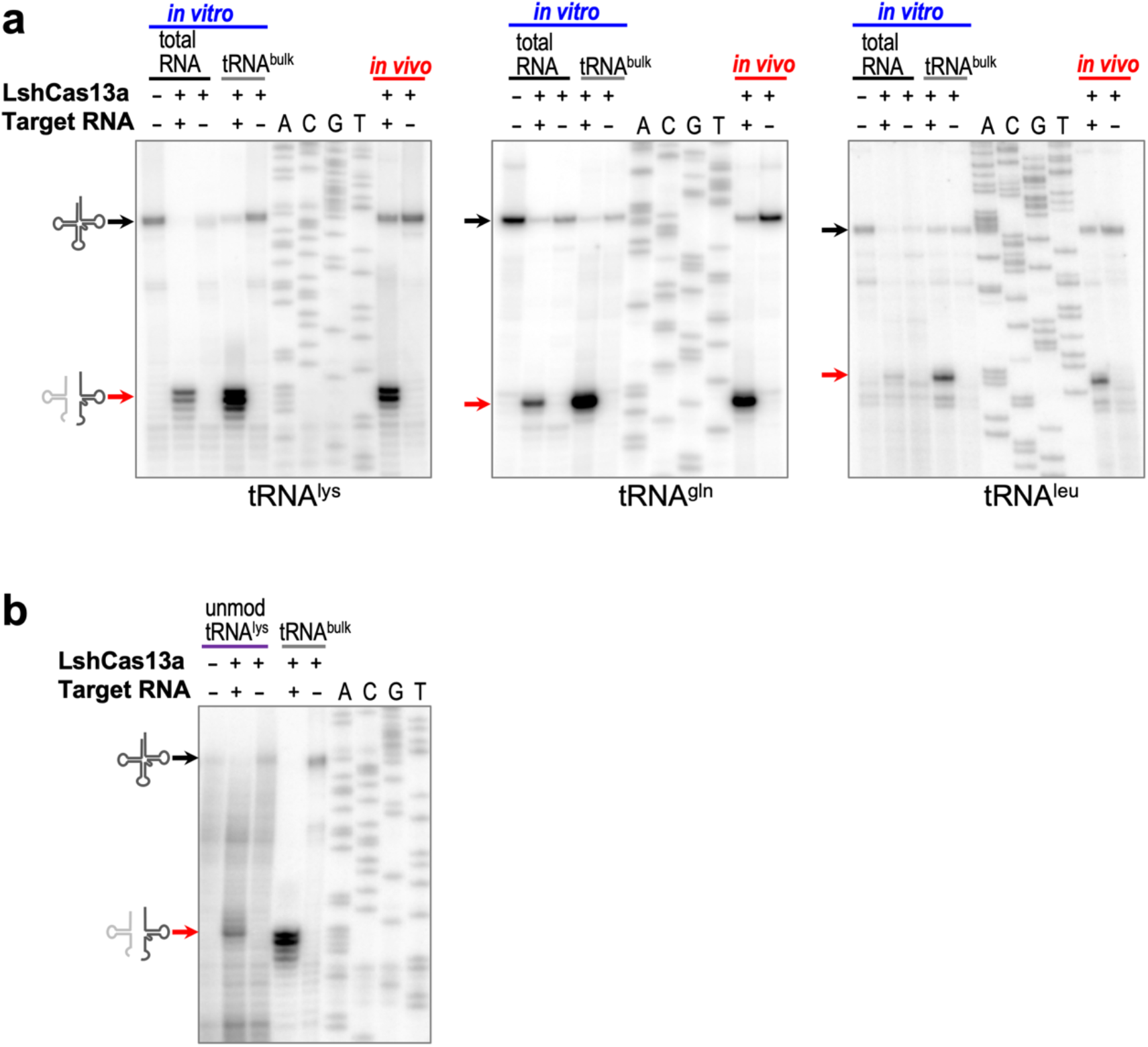
tRNA cleavage *in vitro*. **a**, Total RNA used as a substrate for cleavage by Cas13a *in vitro*. Primer extension assay revealed that position of cleavage sites at tRNA anticodon loop is unchanged when total *E. coli* RNA or bulk *E. coli* tRNA is used as cleavage substrate *in vitro* and matches those observed *in vivo*. **b**, Cas13a cleaves synthetic tRNA^lys^ at the anticodon loop - nucleoside modifications in tRNA are not required for cleavage. All gel images are representative of three independent experiments. For gel source data, see Supplementary Fig. 2.

**Extended Data Fig. 8.**
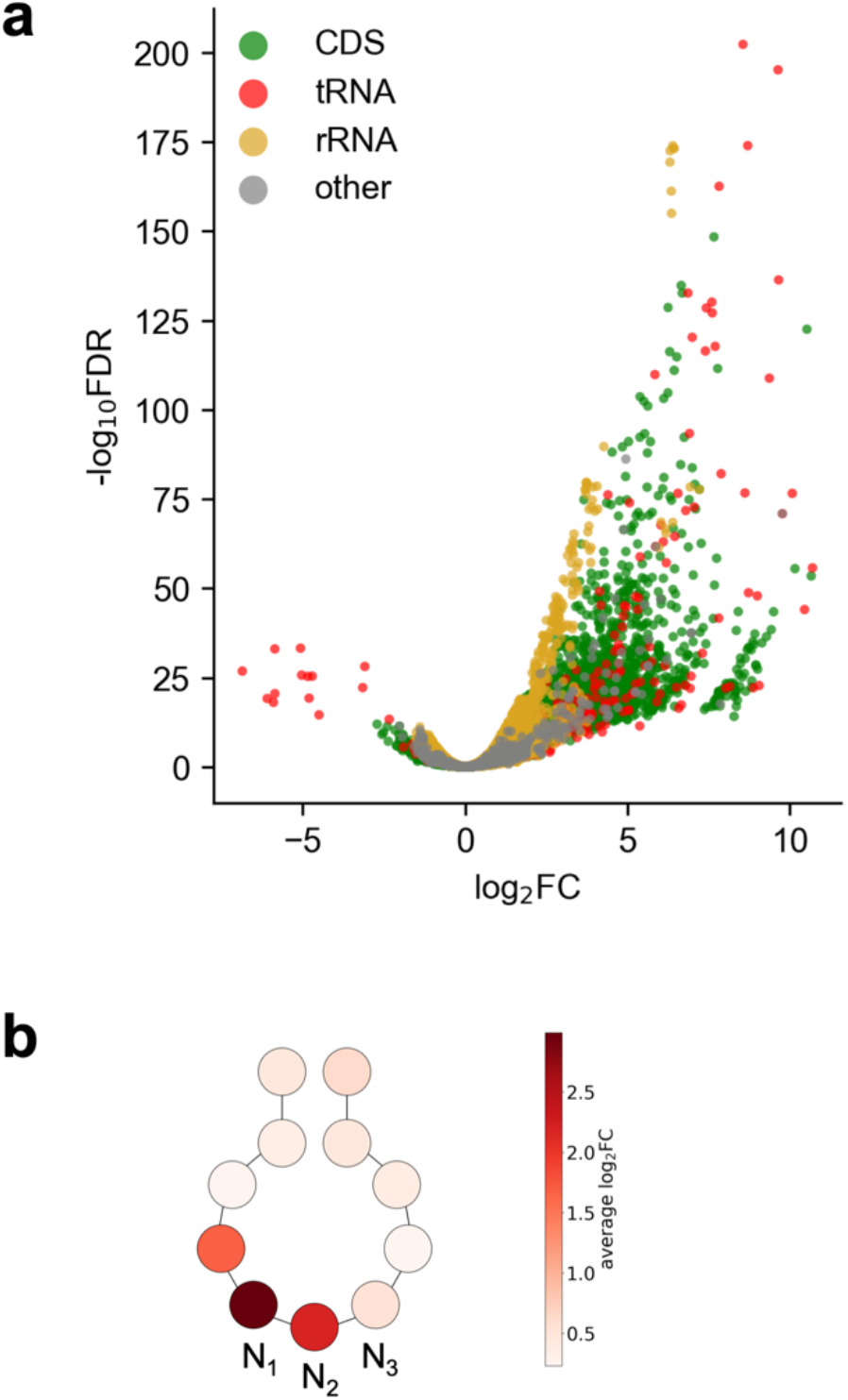
RNA-seq of *in vitro* Cas13a cleavage products. **a**, Volcano plot depicting the results of the comparison of 5’ transcripts ends counts per nucleotide position of each strand between targeting and nontargeting samples in *in vitro* cleavage experiments with isolated total *E. coli* RNA. The analysis was performed in the same way as for Fig. 3d: likelihood ratio test was used. Each dot corresponds to analyzed nucleotide position. The horizontal axis depicts log_2_FC value between targeting and nontargeting samples; the vertical axis depicts adjusted p-value. Dot colors correspond to genomic features where the analyzed nucleotide position is located. **b**, Schematic depiction of tRNA anticodon loop; circles correspond to tRNA nucleotides. Each circle is colored according to color scheme where the intensity of color corresponds to average value of log_2_FC between average 5’ ends CPMs in targeting and nontargeting samples across all tRNA genes detected in *E. coli* genome.

**Extended Data Fig. 9.**
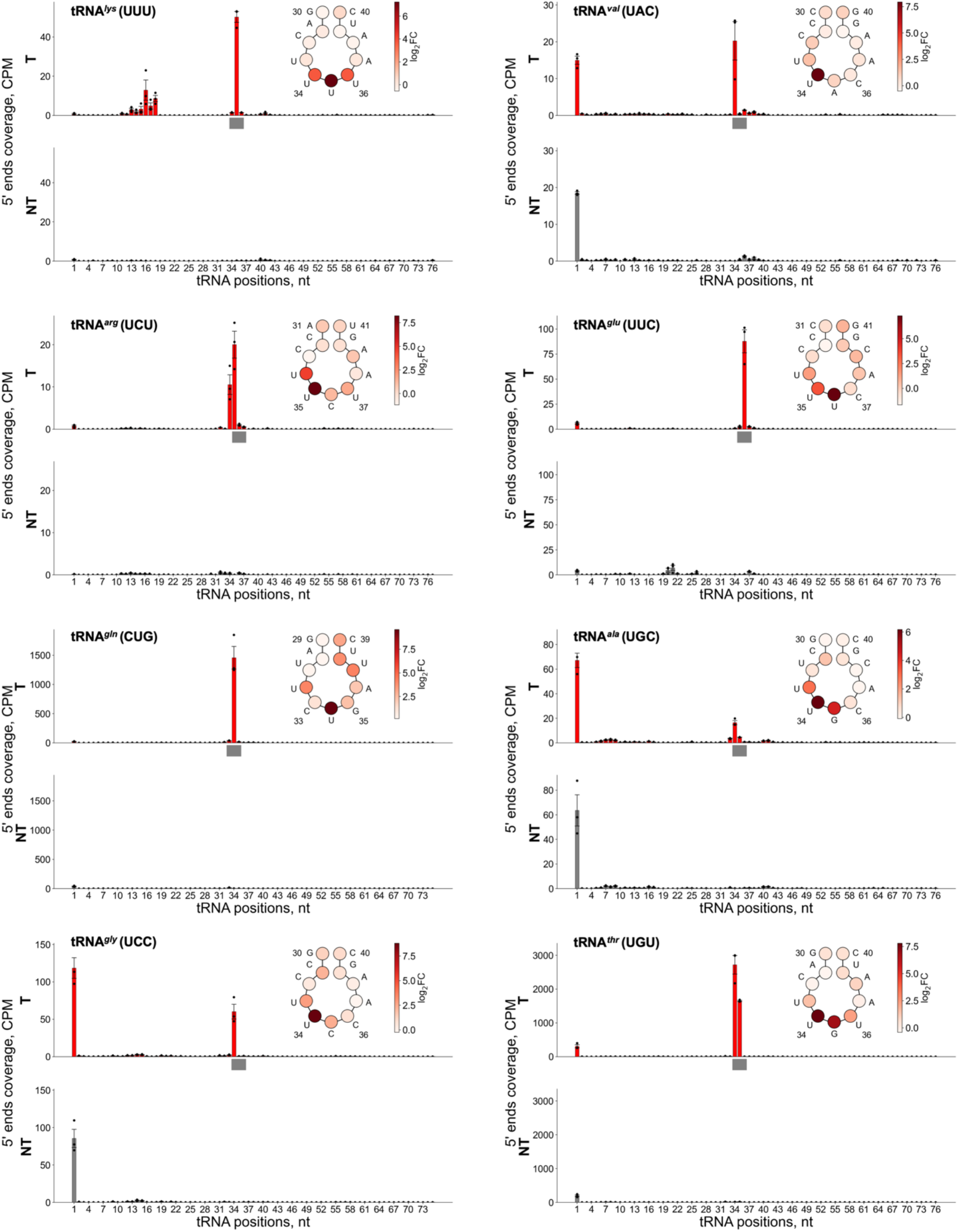
Mapping *in vitro* tRNA cleavage sites. CPM normalized coverage of 5’ ends of transcripts mapped onto tRNA gene in targeting (T, red bars) and nontargeting (NT, grey bars) *in vitro* samples (mean ± s. e. m. from three independent experiments). The horizontal axis depicts the nucleotide position of tRNA. The region corresponding to anticodon is shown by a grey bar. At the top right part of each panel the scheme of tRNA anticodon loop is depicted; circles correspond to tRNA nucleotides. Each circle is colored according to color scheme where the intensity of color corresponds to values of log_2_FC between mean 5’ ends CPMs in targeting and nontargeting samples.

**Extended Data Fig. 10.**
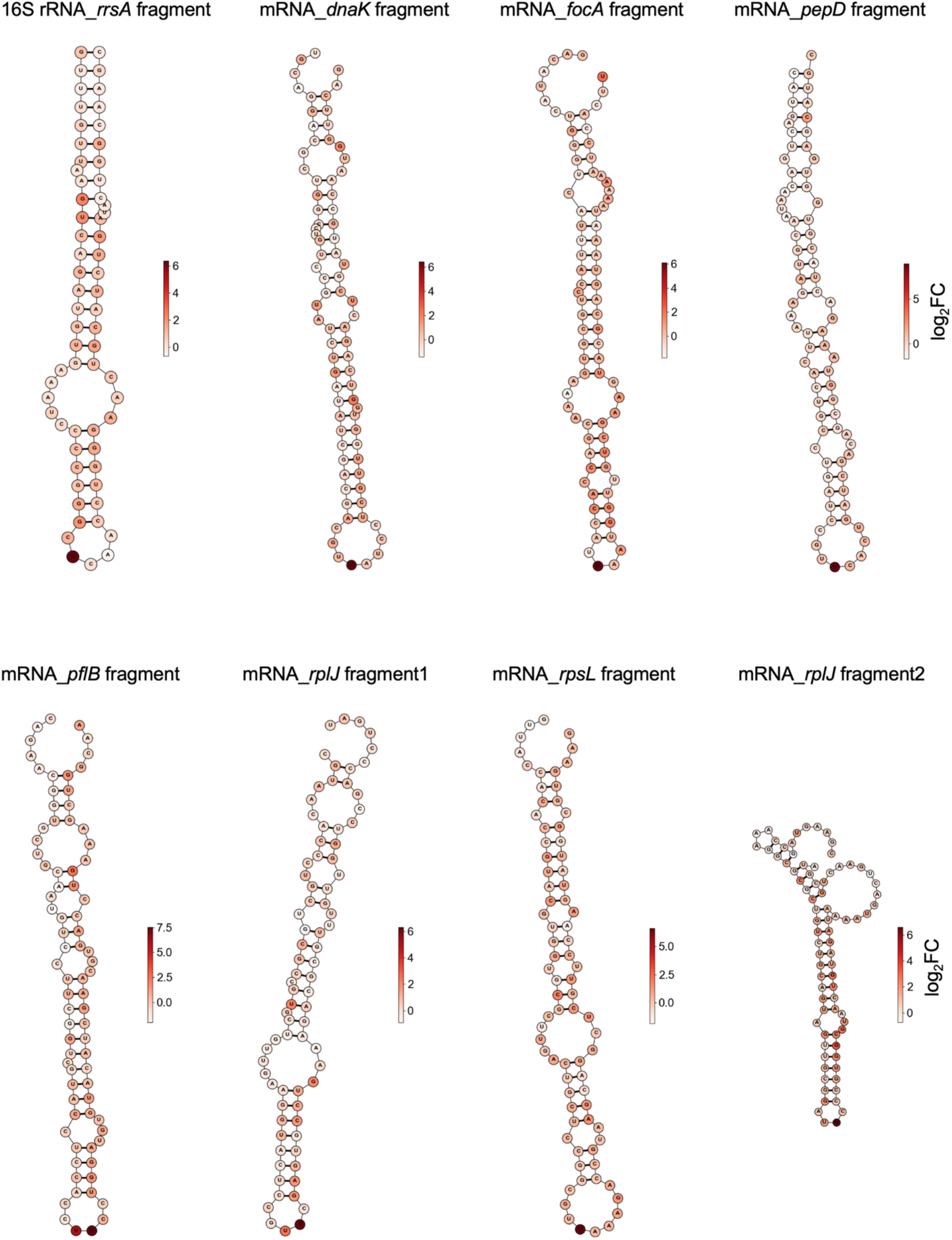
Non-tRNA cleavages by target-activated Cas13a *in vitro.* Mapping of several *in vitro* cleavage sites detected in non-tRNA substrates on the predicted secondary structures of RNA molecules: eighty nucleotides surrounding the analyzed cleavage sites were extracted, and the RNA secondary structures were predicted using RNAfold program (see Methods). Each circle is colored according to color scheme where the intensity of color corresponds to values of log_2_FC between average 5’ ends CPMs in targeting and nontargeting samples. Non-tRNA cleavages at uridine residues located at small loops mimic tRNA cleavages at *in vitro* conditions.

